# Preclinical efficacy of a systemically-administered, second-generation STING agonist that promotes antitumour immunity in combination with radiotherapy

**DOI:** 10.64898/2026.03.23.713628

**Authors:** Malin Pedersen, Lisa Hubbard, Shane Foo, Anton Patrikeev, Antonio Rullan, Holly Baldock, Christian Mandl, Paolo Chetta, Jehanne Hassan, Isaac Dean, Naomi Guppy, Philippe Slos, Charleen Chan, Elizabeth Appleton, Emmanuel C. Patin, Justin Weir, Masahiro Ono, Thorsten Oost, Ulrich Reiser, Thomas Zichner, Kevin Morse, Mary Murphy, Lingqi Luo, Reniqua House, Louise Giffin, Alan Melcher, Anne Vogt, Sebastian Carotta, Kevin Harrington

## Abstract

As potent triggers of innate immunity, STING agonists hold promise as active immunotherapeutic agents for cancer treatment. Second-generation STING agonists, suitable for systemic delivery, are being investigated in preclinical research and have entered clinical trials. Here, the novel synthetic STING agonist, BI-1703880 (STINGa), which was designed for intravenous delivery, was investigated for anti-tumour and immunological effects. We show that STINGa activates the STING pathway and results in a transient and dose-dependent upregulation and secretion of interferons and proinflammatory cytokines *in vitro* and *in vivo*. We show that intravenous administration of repeated dosing with low-dose STINGa is well tolerated. We report that radiotherapy (RT) and STING agonism synergizes to generate innate immune cell and CD8+ T cell responses that control tumour growth. Anti-tumour activity induced by combined RT / STINGa was reduced in mice lacking a functional immune system. RT / STINGa combination treatment also initiated development of protective immune memory. RT / STINGa upregulated PD-L1, PD-1 and CTLA-4 in the tumour microenvironment. Our findings show that combining RT / STINGa with immune checkpoint inhibitors further increases therapeutic benefit. Our data confirm STING as a therapeutic target in cancer and support the clinical development of BI-1703880 STING agonist, thereby suggesting radiotherapy as a potential combination for enhancing anti-tumour efficacy.

## Background

Therapies that (re)invigorate anti-tumour immune responses have transformed cancer care in the last decade. Immune checkpoint inhibitors that target programmed death-1 (PD- 1)/programmed death-1 ligand (PD-L1) and cytotoxic T-lymphocyte-associated protein 4 (CTLA-4) on T, and other, immune cells can induce regressions and even durable complete responses in some cancer patients. However, despite the clinical successes of these therapies, most tumours fail to respond, and this is particularly true for those with tumours devoid of functionally capable tumour-specific CD8+ T cells. For these patient groups, therapeutic strategies capable of inducing *de novo,* or reactivation of existing, anti-tumour immune responses are needed. This requires cancer cell killing with subsequent release of tumour-associated antigens and recruitment and activation of crucial immune cells, such as antigen- presenting cells (APCs). APCs take up antigens released within the TME, mature and migrate and, then, efficiently prime responses by presenting processed forms of the antigens to CD8+ T cells in draining lymph nodes (1).

Radiotherapy is a standard-of-care treatment for many cancers. In addition to its direct cytotoxic effects, radiotherapy can induce immune responses, through micronucleus generation and activation of the cyclic GMP-AMP synthase/stimulator of interferon genes (cGAS/STING) pathway in cancer cells themselves or in neighbouring cells in the TME by transfer of nucleic acids (2). The immunologic effects of radiotherapy critically depend on the cGAS/STING pathway (3, 4).

STING belongs to the family of nucleic acid sensors and is the adaptor for cytosolic DNA signalling. In its basal state, STING exists as a dimer with its N-terminal domain anchored in the endoplasmic reticulum (ER) and its C-terminal domain residing in the cytosol. Cyclic-di- nucleotides (CDNs), generated by cyclic GMP-AMP synthase (cGAS), are the natural ligands of STING (5). Binding of CDNs to STING induces conformational changes which allows the binding and activation of TANK-binding kinase (TBK1) and interferon-regulatory factor 3 (IRF3) and relocation of STING from the ER to the perinuclear endosomes (6). Phosphorylation of the transcription factors IRF3 and NFκB by TBK1 results in expression of multiple cytokines, including type 1 interferons and the activation of APCs. Recent pre-clinical studies have reported that deletion of either cGAS or STING genes in immune cells severely reduces the anti-tumour effects of radiotherapy in tumour-bearing mice, while pathway activation by synthetic STING agonists increases the immunostimulatory effects of radiotherapy (7–9).

First-generation STING agonists were evaluated in clinical trials of direct intratumoural injections based on promising pre-clinical data from mouse models (10). However, systemic delivery of second-generation STING agonists enables simultaneous targeting of multiple tumour lesions and primary / secondary lymphoid tissues at doses that are clinically tolerable. Currently, several systemically-administered synthetic STING agonists are being evaluated in clinical trials.

In this study, we present data for a novel, intravenously-administered STING agonist (STINGa) BI 1703880, that is currently being evaluated in a clinical trial (clinicaltrials.gov NCT05471856). We observed STINGa dose-dependent effects on STING pathway activation and cytokine release *in vitro* and *in vivo*. Our preclinical data suggest that, although efficient for tumour growth inhibition in several mouse tumour models, high-dose, single-agent STINGa is expected to be toxic, and not tolerated, if used repeatedly. We, therefore, hypothesized that a similar anti-tumour effect could be achieved with lower doses of STINGa combined with standard-of-care therapy that induces cancer cell death and stimulates immune responses. Our results show that radiotherapy can successfully be combined with lower, well-tolerated doses of STINGa *in vivo*, resulting in successful initiation of immune-dependent anti-tumour efficacy that is associated with an increase in CD8+ T cells in tumours and upregulation of inhibitory PD-1 and CTLA-4 checkpoints. Consequently, the immune-mediated efficacy of combined STINGa-radiotherapy was further improved by adding anti-PD-1/anti-CTLA-4 immune checkpoint inhibitors. These data provide a rationale for radiotherapy as a potential combination partner.

## Results

### BI-1703880 is a potent activator of the STING pathway

Dendritic cells (DCs) have been proposed as a key target cell population for STING agonists by virtue of their high expression of STING protein and observations that therapy responses are reduced in their absence. Therefore, we used the human monocytic cell line, THP-1 cells, as well as human donor-derived peripheral blood mononuclear cells (PBMCs) and human PBMC- derived dendritic cells for initial studies to assess the specificity of the novel BI 1703880 STINGa and its ability to activate the STING pathway *in vitro*. To test for drug-on-target and compound specificity, we generated paired THP-1 cell lines: parental wildtype (WT) STING or STING-knockout (KO). STING protein knockdown was confirmed in the STING-KO cells (Figure 1A). Then, we tested STING pathway activation and upregulation of downstream events. WT and STING-KO THP-1 cells were treated with STINGa *in vitro* and pathway activation was defined via phosphorylation of STING, TBK1 and IRF3 using Western blot analysis. Strong phosphorylation of the pathway components was observed in STING-WT, but not in STING - KO, cells (Figure 1A). These results were confirmed in primary human PBMCs from three different donors, with STINGa triggering dose-dependent phosphorylation of STING, TBK and IRF3 (Figure 1B).

**Figure 1.**
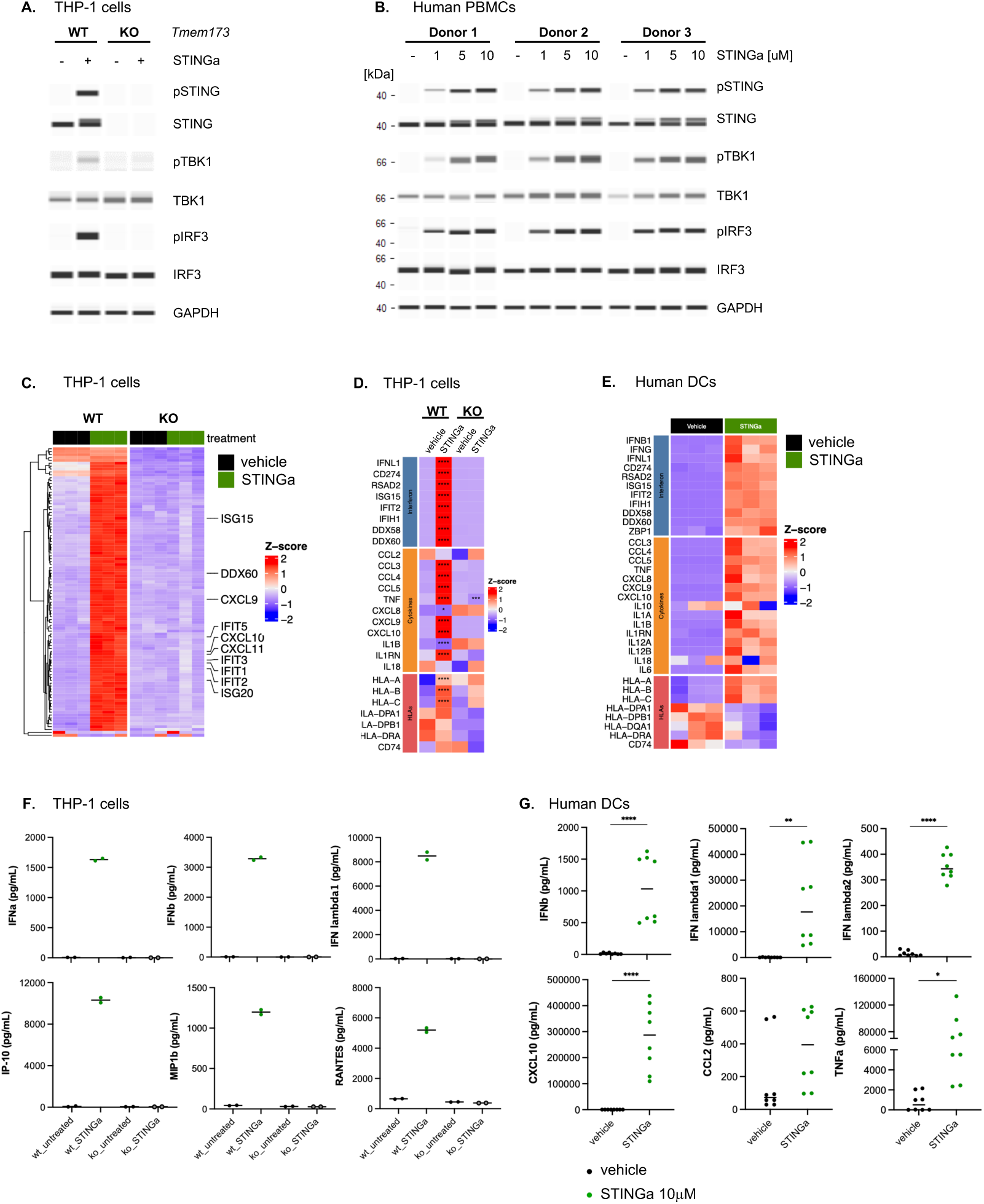
STING agonist activates the STING pathway and results in cytokine secretion and activation of dendritic cells *in vitro* and *in vivo*. **(A)** Wild-type (wt) or STING knockout (ko) (*TMEM173 ko*) THP1 cells were treated with STINGa (10mM, 1 hour) and analysed with Western Blotting using antibodies against phospho-STING, STING, phospho-TBK1, TBK1, phospho-IRF3 and IRF3. GAPDH was used as loading control. **(B)** Human PBMCs from three different donors were treated with a dose-range of STINGa (1, 5, 10mM or left untreated, 30min) and subsequently lysed and analysed for STING pathway activation using Western Blotting and antibodies against phospho-STING, STING, phospho-TBK1, TBK1, phospho-IRF3 and IRF3. GAPDH was used as loading control. **(C)** THP-1 wild-type or STING knockout cells were treated with STINGa for 6 hours *in vitro* and processed for RNA sequencing analysis. Gene expression heat maps separating samples based on top-100 most variable genes. Samples are split by STING knockout (supervised clustering). **(D)** Gene expression analysis from THP-1 cells based on selected interferon genes, cytokines and HLA transcripts. **(E)** Human DCs were treated with STINGa for 6 hours *ex vivo* and processed for RNA sequencing analysis. Gene expression heat maps by sample and by treatment with differential expression (DE) significance are presented. **(F)** Cytokine secretion from wild-type or STING knockout THP1 cells treated with STINGa (5mM or left untreated, 24 hours) were analysed using Bio-Plex assay. **(G)** Cytokine secretion from human dendritic cells (DCs) isolated from PBMCs treated with STINGa *in vitro* (10 mM, 24 hours) were analysed using Bio-Plex assay. Unpaired t-test was used to test for significant differences between groups.

To evaluate further the specificity of STINGa, we examined global transcriptomic changes using RNA sequencing analysis, in both human THP-1 and dendritic cells. STINGa-treated THP-1 cells showed upregulation of immune genes compared to untreated cells, occurring specifically in STING-WT, but not STING-KO, cells (Figures 1C-D). Similar results were obtained from primary human DCs, with a clear induction of interferon-related genes and cytokines following STINGa treatment *ex vivo* (Figure 1E). Notably, we also saw an upregulation of HLA- A, HLA-B and HLA-C genes, known to be critical for cross-presenting antigens to CD8+ T cells (Figures 1D-E). Thereafter, we confirmed functional activation via STING pathway by documenting proinflammatory cytokine release (IFNα, IFNβ, IFNλ, CXCL10, CCL2, CCL5, TNFα) from both THP-1 and dendritic cells following STINGa treatment *in vitro* and *ex vivo* (Figures 1F-G).

Human STING exists in five different variants known as WT (R232), REF (R232H), HAQ (R71H, G230A, R293Q), AQ (G230A, R293Q) and Q (R293Q). We generated THP1-BlueISG-STING reporter cells ectopically expressing the five different variants to determine the activity of BI 1703880 STINGa on each variant. These cells reflect STING-mediated IRF3 activation through the activity of secreted embryonic alkaline phosphatase (SEAP) reporter gene. Importantly, STINGa was active on all five human variants while no reporter signal was detected in THP1 cells lacking STING expression (data not shown).

Together, these results highlight that BI 1703880 STINGa specifically allows for robust STING pathway activation, including upregulation of key immune transcripts and cytokine secretion from human immune cells.

### Systemic treatment with STINGa induces inflammatory cytokines in plasma and tumour tissue and mediates dose-dependent anti-tumour effects

STING activation upregulates cytokines that bridge the innate and adaptive immune systems. Such a cytokine response needs to be finely balanced *in vivo* to receive the benefits, while at the same time reducing unwanted effects of high cytokine levels. Therefore, we examined differential responses and cytokine expression levels of various dose-schedules of intravenously- administered STINGa *in vivo*. For these studies, we used three different murine tumour models, TBPt-4C4 (anaplastic thyroid carcinoma), mEER (head and neck squamous cell carcinoma) and MC38 (colorectal cancer), all implanted subcutaneously into immunocompetent mice.

To determine if intravenous injections of STINGa could induce effective immune responses *in vivo*, we first examined the levels of pro-inflammatory cytokines in plasma and tumour tissue. We assessed STINGa dose-dependency for cytokine secretion by using one high (20 μmol/kg) and one low (7 μmol/kg) intravenous dose in tumour-bearing mice. Following STINGa treatment, we observed drug dose-dependent upregulation of cytokines (IFNα, IFNβ, CCL2, CCL5, IL6, TNFα, CXCL1 and CXCL10) in both plasma and tumour tissue (Figure 2A and Supplementary figure 1A). Single high dose administration of STINGa resulted in very high plasma cytokine levels (ranging from 1000-60000 pg/mL), suggesting that toxicity might occur with repeated administration at high dose. Dosing with low dose also induced cytokines in plasma and tumours, but to lower levels (Figure 2A, Supplementary figure 1A).

**Figure 2.**
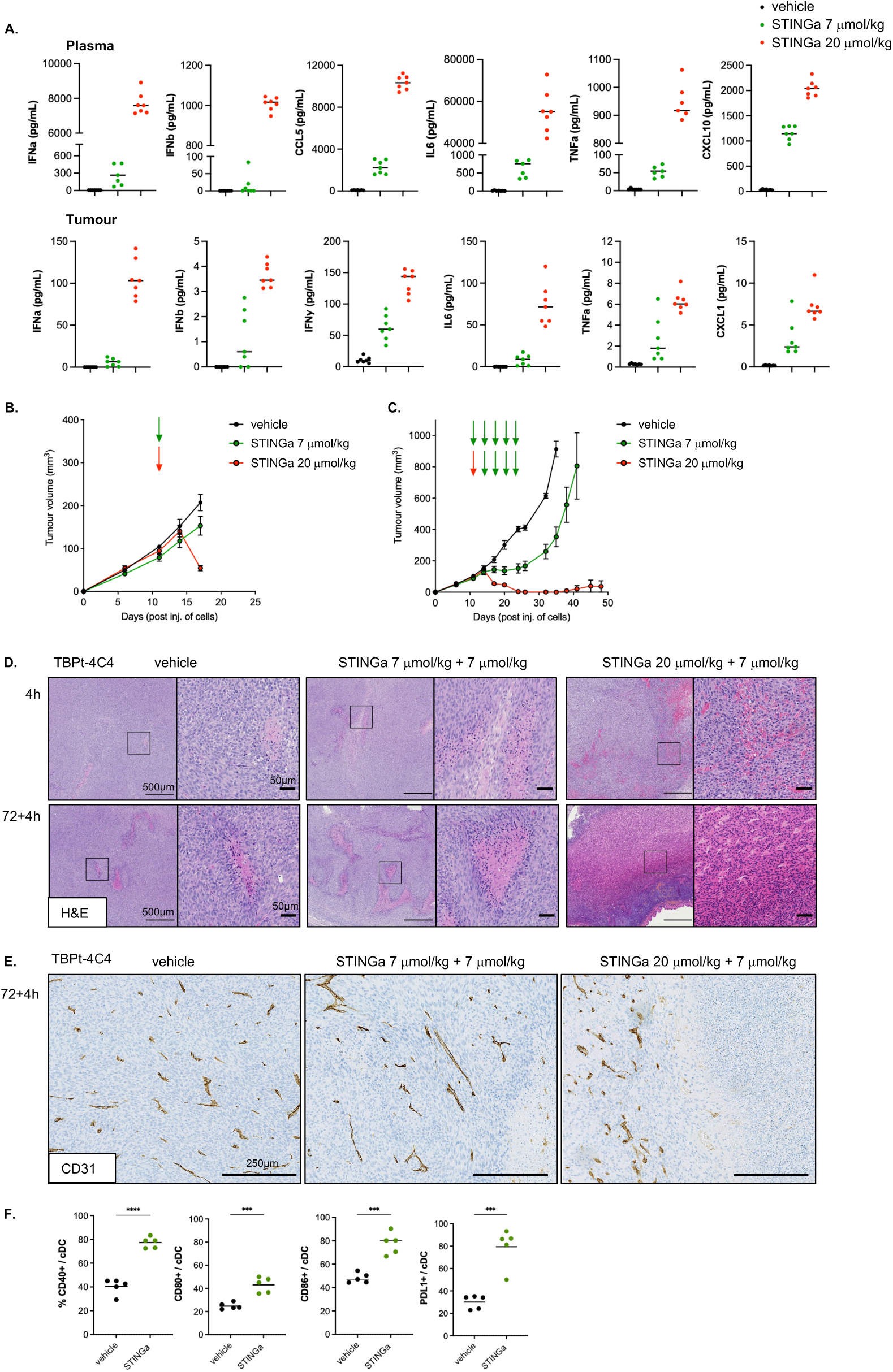
The effect of STING agonist on plasma and tumour cytokine levels, and on tumour growth retardation, is dose- and time-dependent. C57Bl/6 mice were subcutaneously injected with TBPt-4C4 tumour cells and treated by intravenous injections with 7 mmol/kg or 20 mmol/kg of STING agonist. **(A)** Cytokine levels in blood and tumours 4 hours after a single dose of intravenous STING agonist at either 7 mmol/kg or 20 mmol/kg. **(B)** Tumour growth graphs of mice subcutaneously injected with TBPt-4C4 tumour cells and systemically treated with 1 dose of either 7 mmol/kg or 20 mmol/kg of STINGa. **(C)** Tumour growth graphs of C57Bl/6 subcutaneously injected with TBPt-4C4 tumour cells and systemically treated with 5 doses of 7 mmol/kg of STING agonist or 20 mmol/kg for one injection, followed by 4 injections of 7mmol/kg. 1 out of 7 mice were cured of tumour in the STINGa 7mmol/kg (x5 inj) group and 6 out of 7 mice were cured of tumour in the STINGa 20 mmol/kg (x1) + 7 mmol/kg (x4 inj) group. Mice bearing TBPt-4C4 tumours were treated with STING agonist 7 + 7 mmol/kg or 20 + 7 mmol/kg with 72 hours in between the two injections. Tumours were collected 4 hours after the second injection with STING agonist. **(D)** Representative H&E images, scale bars: 500 or 50mm. **(E)** Representative CD31-stained images, scale bars 250mm. **(F)** Tumour-draining lymph nodes (tdLN) from TBPt-4C4 tumour-bearing mice were analysed using flow cytometry 24 hours after the second injection of STING agonist (7 mmol/kg). Percentage of CD40+ / CD80+ / CD86+ or PD-L1+ cDCs were plotted and unpaired t-tests performed to test for significancy between groups.

Next, we investigated the effect of single-agent STINGa therapy on tumour growth using high and low doses. One single intravenous high-dose of STINGa yielded rapid and appreciable tumour shrinkage within 72 hours, while a single low dose had only marginal effects on tumour size (Figure 2B). The rapid tumour regression following high-dose STINGa injection suggested that non-immune-related effects were occurring. Therefore, we analysed tumours using immunohistochemistry and compared the short-term effects of injections of either high- or low-dose STINGa. At the initial time-point (day 14 after subcutaneous cell injection), there are limited areas of necrosis in the TBPt-4C4 tumours treated with vehicle. Histological analysis of hematoxylin & eosin (H&E)-stained tumour sections revealed large areas of tumour necrosis following high-dose STINGa, with visible haemorrhage, red blood cell extravasation and neutrophil infiltration. In contrast, low-dose STINGa did not lead to a meaningful increase of necrotic areas, although we could observe perivascular necrotic foci. Comparing one vs two injections of STINGa revealed that STINGa treatment-induced tumour cell death in an apparent dose- and time- dependent manner (Figure 2D). Single-dose intravenous STINGa injection had a similar effect on mEER tumours. However, in this model, significant necrosis was also observed following low-dose STINGa treatment, whereas high-dose administration induced extensive central tumour cell death, leaving a surrounding rim of viable cells (Supplementary Figure 1G). CD31 immunostaining revealed a less complex vascular network following low-dose STINGa and especially in contrast to high-dose STINGa, with absence of blood vessels at the necrotic areas (Figure 2E and Supplementary figure 1G). Notably, vessels outside the tumours remained unaffected.

To investigate the long-term effects of different doses of single-agent STINGa on tumour growth, we administered STINGa systemically twice-weekly for a total of 5 doses to TBPt-4C4 tumour-bearing mice. Following the observed high-level cytokine secretions after high-dose STINGa, we compared one single high dose followed by four low doses versus five low doses. The repeated low-dose, single-agent STINGa showed moderate tumour growth control, while one high-dose of STINGa followed by 4 low doses eradicated 6 out of 7 tumours in line with the massive necrosis seen following a single injection of high-dose STINGa (Figure 2C). We did not detect any significant long-term weight loss in mice treated with repeated low dosing of STINGa and selected this dose-schedule for further experiments (Supplementary figure 2M). We assessed cytokine levels following low-dose STINGa in a time-course and found time- dependent differences in the secretion of various cytokines. In both plasma and tumour, the earliest cytokines (IFNα, IFNβ, TNFα) peaked as early as 1 hour after STINGa injection, while others (CCL2, CCL5, IFNγ) peaked at 4 hours. All cytokines investigated in plasma had returned to baseline levels by 24 hours (Supplementary figure 1B). In tumours, type I interferons rapidly returned to pre-treatment levels, but IFNγ, and chemokines CCL2 and CCL5, were still upregulated at 24 hours (Supplementary figures 1B-C). Since IFNβ is a major downstream target of STING, we performed *in situ* hybridisation for IFNβ in MC38 tumours (Supplementary figure 1D). This assay confirmed early upregulation of IFNβ in tumours and validated that induction of IFNβ was both time- and STINGa dose-dependent.

Dendritic cells are STINGa targets, key mediators of immunity and known to be required for effective T cell activity against tumours. Therefore, we investigated if intravenous, low-dose STINGa could activate this cell population *in vivo*. We detected increased expression of activation markers (CD40+, CD80+, CD86+, PD-L1) on murine conventional dendritic cells (cDCs) in lymph nodes from tumour-bearing mice from all three models following systemic STINGa administration *in vivo* (Figure 2F and Supplementary figures 1E-F). Furthermore, single agent STINGa treatment *in vivo* upregulated Ifng, PD-L1 and PD-L2 transcripts as well as protein levels of PD-L1 in tumours (Supplementary figures 1K-L).

Although we have not fully assessed the relative contributions of STING activation within tumour versus host cells in our models, Western blot analysis of TBPt-4C4, mEER and MC38 cells treated with STINGa *in vitro* revealed that TBPt-4C4 cells express STING protein and activate the STING-pathway, while mEER and MC38 tumour cells do not (Supplementary figures 1H-I).

Taken together, these results show a rapid, but transient, dose-dependent release of cytokines in the circulating blood and tumour microenvironment following systemic STINGa injection. Our data suggest that multiple, repeated low-dose STINGa injections might boost immune activation and, in concert with conventional cancer treatment modalities, exert synergistic anti-tumour effects.

### The combination of radiotherapy and STING agonism improves therapeutic outcome and is dependent on a functional immune system

Building on the promise of using repeated intravenous dosing of 7 μmol/kg STINGa therapy to enhance immune activation, we hypothesised that standard-of-care radiotherapy (RT) might be an effective combination partner with STINGa through direct cytotoxicity and increased release of tumour-associated antigens. To explore this, we first assessed TBPt-4C4 and mEER tumours for radiosensitivity and treatment-induced infiltration of CD8+ T cells. Two fractions of 8Gy RT on consecutive days resulted in tumour growth retardation and increased T cell abundance in the TME (Supplementary figures 2A-E). Then, we investigated the effects of RT on MHC-I and II on tumours using bulk RNA sequencing and observed an increase in HLA transcripts following RT (Supplementary figure 2E). An increased expression of MHC-I and PD- L1 on tumour cell lines treated with RT *in vitro* was also observed (Supplementary figure 2F). Taken together, these data suggest that RT can retard tumour growth and increase presentation of antigens by tumour cells for targeting by cytotoxic T cells.

Next, we tested the therapeutic efficacy of low-dose systemic STINGa added as an immune booster to initial RT (two fractions of 8Gy on consecutive days). Tumours were monitored for growth, and survival was based on tumour burden (Figure 3A). Mice were weighed twice-weekly and exhibited transient weight loss following radiotherapy (Supplementary figures 2P-R). Mice bearing TBPt-4C4 or mEER tumours showed significantly improved tumour control and survival with combined RT/STINGa over single-agent therapies or vehicle-treated controls (Figures 3B-C). Mice that achieved treatment-induced tumour cure received a rechallenge injection on the contralateral flank. Importantly, this showed development of protective immune memory, suggesting involvement of the adaptive immune system (Figures 3B-C). Thereafter, we delayed the onset of treatment of TBPt-4C4 tumours until the tumours were larger in size. This still resulted in significant synergistic effect of combination therapy, although none of the tumours was cured (Supplementary figure 2G). Improved tumour growth control with combined RT and systemic STINGa was subsequently confirmed in five additional subcutaneous mouse tumour models (MC38, TC1, AT84, TBP-B79 and TKP-4831) (Supplementary figures 2H-I) and in mEER tumours growing orthotopically in the lip (Supplementary figure 2J).

**Figure 3.**
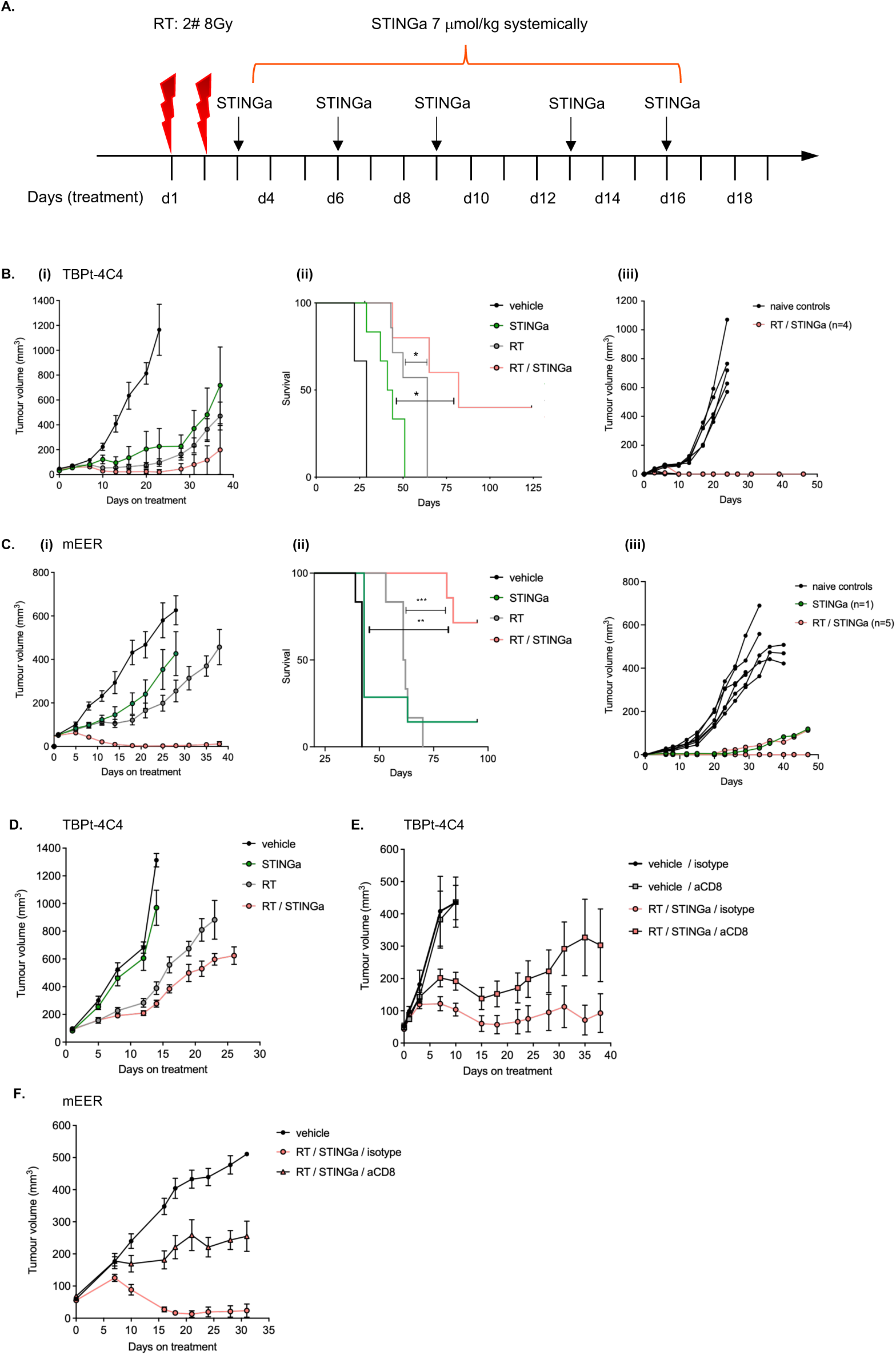
The combination of radiotherapy and STING agonist improves therapeutic outcome and is dependent on a functional immune system. **(A)** Schematic of the treatment schedule. Wild-type C57Bl/6 or NOD scid gamma (NSG) mice were subcutaneously injected with tumour cells as indicated and treated with 2 fractions (#) of 8Gy of radiotherapy on consecutive days followed by 5-6 systemic injections (inj.) of STING agonist at 7mmol/kg, every 3-4 days. Vehicle solution was given at the same days as STING agonist to all mice in vehicle or RT cohorts. **(B)** Tumour growth **(i)**, survival **(ii)** and rechallenge **(iii)** graphs of mice treated with radiotherapy and STING agonist (6 inj.) in the TBPt-4C4 subcutaneous model, n=8 mice / group. Log-rank (Mantel-Cox) test was performed on Kaplan-Meier survival graphs: vehicle vs RT/STINGa: P value 0.0014 (**), RT vs RT/STINGa: P value 0.0241 (*), STINGa vs RT/STINGa: P value 0.0116 (*). **(C)** Tumour growth **(i)**, survival **(ii)** and rechallenge **(iii)** graphs of mice treated with radiotherapy and STING agonist (5 inj.) in the mEER subcutaneous model, n=8 mice / group. Log-rank (Mantel-Cox) test was performed on Kaplan-Meier survival graphs: vehicle vs RT/STINGa: P value 0.0006 (***), RT vs RT/STINGa: P value 0.0002 (***), STINGa vs RT/STINGa: P value 0.0076 (**). **(D)** Tumour growth graphs of NSG mice treated with radiotherapy and STING agonist (6 inj.), n=5-7 mice / group. **(E)** Tumour growth graph of TBPt-4C4 tumour- bearing mice treated with radiotherapy and STING agonist (5 inj.) and an antibody for CD8 depletion. **(F)** Tumour growth graph of mEER tumour-bearing mice treated with radiotherapy and STING agonist (5 inj.) and an antibody for CD8 depletion.

To investigate the contribution of the immune system to therapeutic efficacy, we used immunodeficient NOD scid gamma (NSG) mice bearing TBPt-4C4 tumours and repeated the therapeutic experiment. Reduced tumour control with combination treatment was observed in NSG mice compared to wild-type mice (Figure 3D and Supplementary figure 2G), confirming the importance of the immune response in therapy. In addition, in TBPt-4C4 and mEER tumour-bearing wild-type mice whose CD8+ T cells were specifically antibody-depleted, the effect of RT/STINGa on tumour growth control was reduced compared to mice with an intact immune system (Figures 3E-F). Efficient depletion of CD8+ T cells in blood and tumour tissue was confirmed by flow cytometry and immunohistochemistry, respectively (Supplementary figures 2M-N). Our observations suggest that CD8+ T cells are central to the efficacy of treatment. Overall, these data show that combined RT/STINGa therapy controlled tumour growth and prolonged survival in multiple murine tumours models in an immune-dependent manner.

### The combination of radiotherapy and STING agonism increases activated CD8+ T cells in tumours

Considering the reduced anti-tumour efficacy of RT/STINGa in CD8+ T cell-depleted mice, we sought to characterise this cell population in tumours. First, we assessed CD8+ T cell infiltration into tumours post-treatment and detected a significant increase in T cells numbers using immunohistochemistry in all four tumour models (TBPt-4C4, mEER, TBK-4831 and MC38) following RT/STINGa combination treatment (Figure 4A and Supplementary figure 3A). To study differences in the TME composition, post-treatment analyses of tumours were performed 10, 12, 14 or 21 days after initiation of RT (different time-points depending on tumour model and assay used for analysis). We characterised the TME using bulk RNA sequencing of TBPt-4C4 tumours to evaluate further the immunological mechanisms underlying regression of tumours. This analysis confirmed the increase of CD3+ lymphocytes in the tumours, including enhanced transcript levels of CD8. In addition, this analysis showed upregulation of transcripts involved in interferon signalling, cytokine/chemokine production, infiltration of various subsets of immune cell and an increase in activation/exhaustion markers for immune cells (Figure 4B). Of interest, the druggable PD-1/PD-L1 and CTLA-4/CD28 immune axes were upregulated following RT/STINGa combination therapy. We confirmed these results in the MC38 model, focusing the transcriptional analysis on hallmark immune pathways (Supplementary figure 3C). Importantly, although the transcriptional analysis also showed increases in CD4+ and FoxP3+ T cells, the ratio of CD8+ T cells/T-regulatory suppressor cells (CD4+/FoxP3+) was increased in both TBPt-4C4 and mEER tumours following combination treatment when analysed on flow cytometry (Figure 4C). Upregulated immune checkpoints and a significant increase of PD-L1 was subsequently validated at the protein level using flow cytometry (Figure 4F, 4H-I and Supplementary figure 3B). Taken together, these results support the notion that RT/STINGa combination treatment creates a more inflamed immune microenvironment in tumours compared to single-agent therapies.

**Figure 4.**
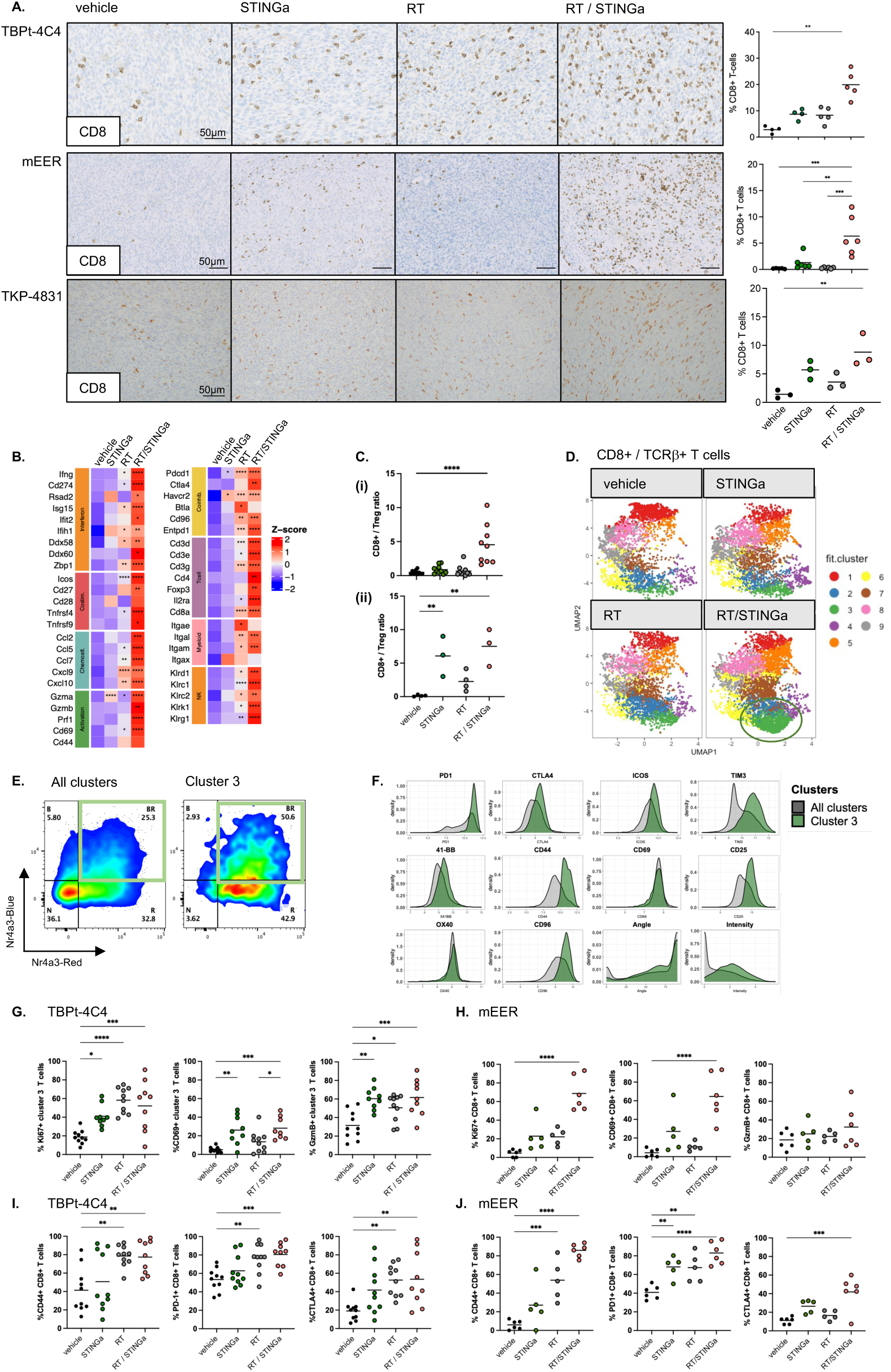
The combination of radiotherapy and STING agonist increases activated CD8+ T cells in tumours. **(A)** Immunohistochemistry images of representative TBPt-4C4, mEER and TKP-4831 tumours stained for CD8 following treatment with radiotherapy (RT) and STING agonist. Tumours were collected 14 days (TBPt-4C4), 13 days (mEER) and 21 days (TKP-4831) after start of RT. Quantification of CD8+ T cells in the tumours using QuPath and graphs plotted to the right. Scale bar 50mm. **(B)** Mice bearing TBPt-4C4 tumours were treated as in Figure 1A and bulk RNAseq was performed on tumours 21 days after start of radiotherapy (n= 3 (vehicle), 3 (STINGa), 5 (RT) and 5 (RT/STINGa) tumours / group). DE significance symbols for adjusted p-values are marked in the heatmaps and each column is compared to vehicle. **(C)** CD8 / Treg (CD4+ FoxP3+) ratios in TBPt-4C4 **(i)** and mEER **(ii)** tumours treated with RT and STING agonist, analysed by flow cytometry. **(D)** UMAP analysis of flow cytometry data of CD8+ / TCR+ T cells from TBPt-4C4 tumour bearing Nr4a3-tocky mice treated with radiotherapy and STING agonist (n=3-5 mice / group), showing cell clusters based on expression markers. **(E)** Representative flow cytometry plots from the experiment in (D) for CD8+ T cells showing Timer Blue and Red fluorescence in all clusters and in Cluster 3. **(F)** Overlayed histograms showing the indicated marker expression on CD8+ T cells in Cluster 3 compared to all concatenated clusters. **(G)** TBPt-4C4 tumour-bearing mice were treated as in Figure 1A and Ki67, CD69 and GzmB expression on “cluster 3” CD8+ T cells in tumours were analysed using flow cytometry 14 days after start of radiotherapy. Data from two experiments were pooled. **(H)** mEER-tumour bearing mice were treated as in Figure 1A and Ki67, CD69 and GzmB expression on CD8+ T cells in tumours were analysed using flow cytometry 10 days after start of RT. **(I)** CD44, PD-1 and CTLA-4 expressing CD8+ T cells in tumours (as in Figure 4G) were analysed using flow cytometry. Data from two experiments were pooled. **(J)** CD44, PD-1 and CTLA-4 expressing CD8+ T cells in tumours (as in Figure 4H) were analysed using flow cytometry. One way ANOVA was performed to test for statistically significant differences between groups.

Next, we used the Nr4a3-timer ‘Tocky’ transgenic mouse model, which allows interrogation of the temporal dynamics of cognate antigen recognition events (11–13), to investigate the effects of RT/STINGa on the temporal dynamics of TCR signalling and to identify markers of highly activated subpopulations of CD8+ T cells. We hypothesised that persistent exposure of CD8+ T cells to tumour antigen will result in an exhausted, dysfunctional phenotype that might be reversed with immune checkpoint inhibition. To test this, TBPt-4C4 tumour-bearing Nr4a3- timer mice were treated with RT and STINGa and the intratumoural CD8+ T cells were analysed by spectral flow cytometry. Following data normalisation, dimension reduction via UMAP and unsupervised clustering of extracellular flow cytometry markers, we first identified a distinct CD8+ T cell cluster (“cluster 3”) that was increased in RT/STINGa-treated tumours compared to vehicle or single agent controls (Figure 4D). We then focused on the cells undergoing persistent TCR-signalling (i.e. Blue+ Red+ CD8+ T cells), and found that the frequency of cells undergoing persistent cognate antigen signalling within “cluster 3” was 50.3% compared to 25.3% in all other clusters taken together (Figure 4E). Moreover, these “cluster 3” CD8+ T cells, persistently signalling through their TCR, expressed higher levels of markers PD-1, CD44, TIM3 and ICOS (Figure 4F). We subsequently further validated the flow cytometry analysis in wild-type mice bearing TBPt-4C4 tumours and by gating for “cluster 3” CD8+ T cells (PD- 1^hi^CD44^hi^TIM3^hi^ICOS^hi^) we confirmed that these cells are also highly proliferative (Ki67+), activated (CD69+) and cytotoxic (GzmB+) (Figure 4G). Activated and proliferating CD8+ T cells were also increased following RT/STINGa treatment in mEER and MC38 tumours, further supporting our results (Figure 4H and Supplementary figure 3D). Using both TBPt-4C4 and mEER tumours models, we thereafter validated PD-1 and CTLA-4 as potential targets for immune checkpoint inhibition to further increase and sustain T cell activation and retard tumour growth (Figures 4I-J). Overall, tumours treated with STINGa and RT/STINGa induced higher expression of activation/exhaustion markers in cells that actively interact with their cognate antigen in a persistent manner. We also analysed CD8+ T cells within tumour-draining lymph nodes following therapy in the TBPt-4C4 and mEER models and observed a similar increase in activation (CD69+) of these cells in a STING-dependent manner (Supplementary figures 3E-F).

To more widely investigate if locally-delivered radiotherapy contributes to beneficial systemic immune effects of STINGa treatment, we set up a bilateral flank tumour experiment, in which the left flank tumour cells were injected 3 days later than the right flank tumours. We locally irradiated only the right flank tumours, while STINGa was delivered systemically. The irradiated, right-sided tumours showed synergistic effects on tumour growth retardation consistent with previous experiments. In the RT single-agent group, we did not observe any abscopal effects on the unirradiated, left-sided tumour following irradiation of the right flank tumours. The effects of single-agent STINGa were greater on the unirradiated tumours, likely due to the smaller size of these tumours at treatment initiation (Supplementary figure 3G). Although we did not observe a significant reduction in unirradiated tumour volume between STINGa- and RT/STINGa-treated animals, there was an increase in CD8+ T cell infiltration in unirradiated, left-sided tumours following RT/STINGa, indicating that the immune response in the unirradiated tumours had benefited from systemic responses that had been initiated by irradiation of contralateral disease (Supplementary figure 3H), albeit not to an extent sufficient to impact on tumour growth.

### The addition of immune checkpoint inhibitors aPD-1 / aCTLA-4 improves therapeutic outcome

Following the observed upregulation of PD-1/PD-L1 and CTLA-4 immune axes in RT/STINGa- treated tumours at both RNA and protein levels, and because these are clinically validated druggable targets, we proceeded to assess if addition of aPD-1/aCTLA-4 to the RT/STINGa treatment *in vivo* would yield increased therapeutic benefit. For these experiments, we used well-established, larger-sized TBPt-4C4 and mEER tumours, that we had previously shown not to be curable by combination RT/STINGa only. Quadruple combination treatment of mice bearing either TBPt-4C4 or mEER tumours resulted in tumour clearance with subsequent protection from a rechallenge injection (5/6 mice cured with quadruple therapy compared to 1/6 in the RT/STINGa/isotypes group in the TBPt-4C4 model and 6/6 mice with quadruple therapy compared to 5/8 in the RT/STINGa/isotypes group in the mEER model) (Figure 5A and Supplementary figure 4A). The addition of both checkpoint inhibitors (aPD-1 and aCTLA-4) was required for maximal effect (Supplementary figures 4B-C). The enhanced therapeutic benefit with quadruple therapy was also reflected in survival curves (Figures 5A, Supplementary figures 4A-D). In a further model, TKP-4831, we detected improved therapeutic benefit in terms of tumour control with the addition of aPD-1/aCTLA-4 to RT/STINGa treatment but without achieving tumour cure. However, the Kaplan-Meier survival curve showed significantly prolonged survival of mice when treated with the quadruple therapy compared to control or those treated with RT/STINGa/isotypes (Supplementary figure 4E).

**Figure 5.**
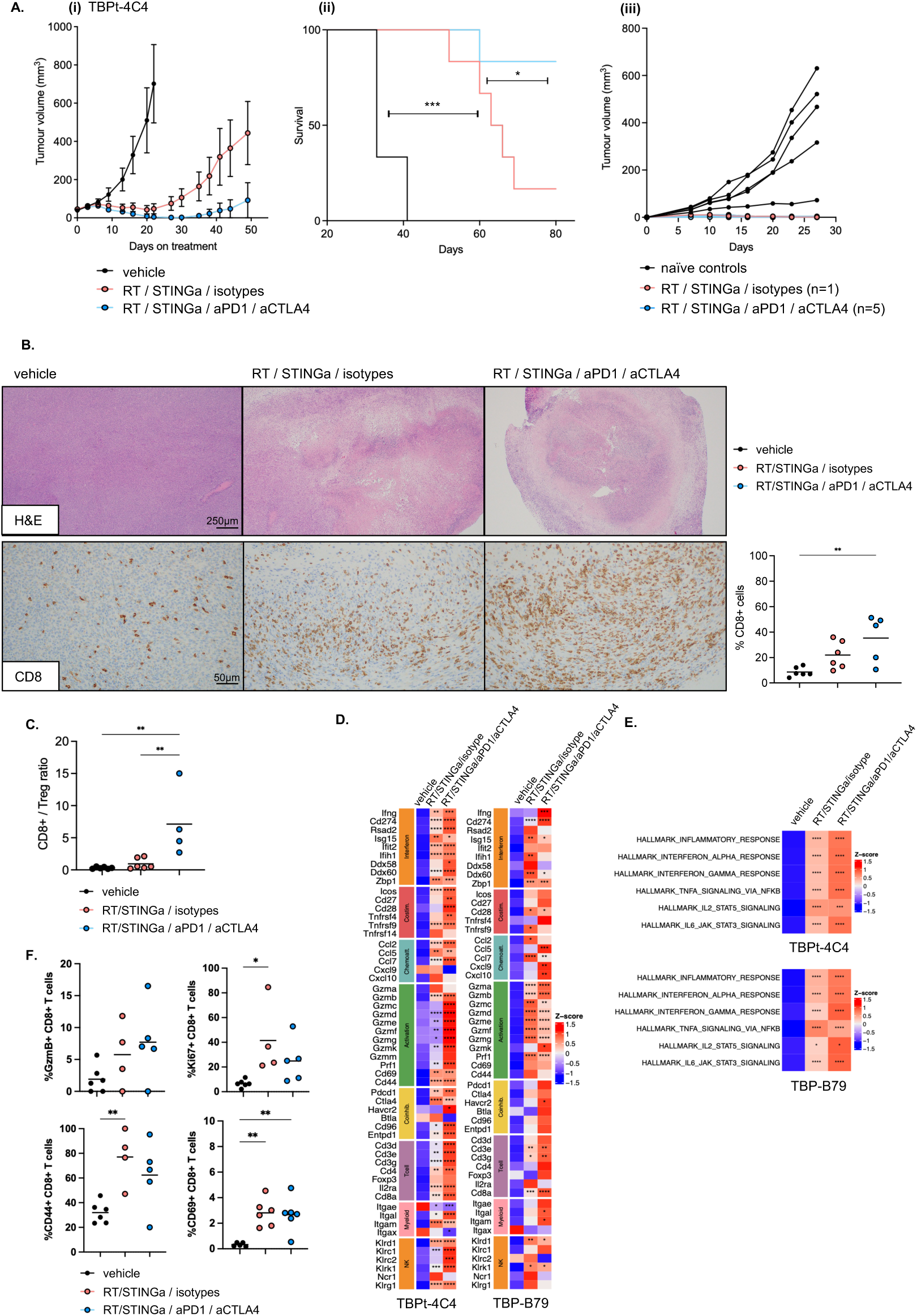
The addition of immune checkpoint inhibitors aPD-1 / aCTLA-4 improves therapeutic outcome. **(A)** Tumour growth **(i)**, survival **(ii)** and rechallenge **(iii)** graphs of mice treated with radiotherapy (2 fractions of 8Gy), STING agonist (5 injections, every 3-4 days) and the combination with immune checkpoint inhibitors aPD1 / aCTLA4, in the TBPt-4C4 subcutaneous model, n=6-7 mice / group. Mice were treated as in Figure 1A with the addition of aPD1 / aCTLA4 on the same days as STING agonist. 1 out of 6 mice were cured of tumour in the RT / STINGa / isotype group and 5 out of 6 mice were cured of tumour in the RT / STINGa / aPD-1 / aCTLA-4 group. Log-rank (Mantel-Cox) test was performed on Kaplan-Meier survival graphs: vehicle vs RT/STINGa/isotypes: P value 0.0006 (***), vehicle vs RT/STINGa/aPD-1/aCTLA-4: P value 0.0006 (***), RT/STINGa/isotypes vs RT/STINGa/aPD- 1/aCTLA-4: P value 0.0345 (*). **(B)** Immunohistochemistry images of representative TBPt-4C4 tumours stained for CD8 following treatment with radiotherapy, STING agonist and aPD-1 / aCTLA-4. Tumours were collected 12 days after start of RT. Quantification of CD8+ T cells in the tumours using QuPath and plotted to the right. **(C)** CD8+ / Treg (CD4+ FoxP3+) ratios in TBPt-4C4 tumours treated with radiotherapy, STING agonist and aPD-1 / aCTLA-4, analysed by flow cytometry 14 days after start of RT. **(D)** Mice bearing TBPt-4C4 or TBP-B79 tumours were treated as in Figure 1A with the addition of aPD-1 / aCTLA-4 on the same days as STING agonist, and bulk RNAseq was performed on tumours 14 days after start of radiotherapy and analysed for single transcripts or **(E)** using hallmark pathway analysis. DE significance symbols for adjusted p-values are marked in the heatmaps, all comparisons are versus vehicle. TBPt- 4C4 tumours: n= 8 (vehicle), 7 (RT/STINGa/isotypes) and 7 (RT/STINGa/aPD1/aCTLA4) and (TBP-B79 tumours: n= 4 (vehicle), 5 (RT/STINGa/isotypes) and 5 (RT/STINGa/aPD1/aCTLA4). (F) TBPt-4C4 tumours-bearing mice were treated as in (C) and CD8+ T cells in tumours were analysed using flow cytometry. One way ANOVA was performed to test for statistically significant differences between treatment groups.

Subsequently, we used immunohistochemistry to analyse TBPt-4C4 tumours two weeks after initiating treatment with RT/STINGa and immune checkpoint inhibitors. Vehicle-treated tumours contained no signs of necrosis, fibrosis or inflammation. Tumours treated with RT/STINGa/isotypes or RT/STINGa/aPD-1/aCTLA-4 exhibited extensive central necrosis and fibrosis, accompanied by active chronic inflammation, with only limited residual viable tumour (Figure 5B). Although H&E images did not reveal obvious differences in cell shape or patterns with the addition of checkpoint inhibitors, larger areas of necrosis were seen following quadruple therapy, and two tumours had no remaining viable tumour (Figure 5B). Furthermore, CD8-specific staining revealed increased infiltration of CD8+ T cells into TBPt- 4C4 tumours upon the addition of aPD-1/aCTLA-4 which was also reflected in the increased CD8+/Treg ratio observed using flow cytometry (Figure 5B and 5C). Bulk RNA sequencing of tumours demonstrated that the transcriptional profiles of tumours treated with RT/STINGa/isotypes vs RT/STINGa/aPD-1/aCTLA-4 were similar, with increases in granzymes and perforin transcripts being the most profound differences with the quadruple therapy. RNA transcript levels also confirmed an increase in T cell infiltration in tumours treated with the quadruple therapy (Figure 5D). Furthermore, to gain a broader insight into immune modulation within TBPt-4C4 and TBP-B79 tumours, we analysed transcriptional data using hallmark pathways and observed a significant increase in inflammatory response pathways with the addition of aPD-1/aCTLA-4 to the backbone RT/STINGa therapy and an increase in interferon alpha and gamma pathways, although this did not reach significance (Figure 5E). By traditional flow cytometry, we did not detect any significant differences in proliferation (Ki67+) or activation marker expression (CD69+, CD44+) on CD8+ T cells within the tumours treated with double vs quadruple combination therapies (Figure 5F). Therefore, we posit that the addition of aPD-1/aCTLA-4 further activates type I and II IFN responses in the TME and promotes tumour clearance by the immune system. Hence, even though there are greater numbers of CD8+ T cells in the RT/STINGa/aPD-1/aCTLA-4-treated animals, they are phenotypically very similar to those seen with RT/STINGa/isotypes but less exhausted and capable of greater effector function (e.g. granzyme release) in the presence of aPD-1/aCTLA- 4 blockade. Therapeutically, this yields the observed benefits in tumour growth retardation, increased cure rates and prolonged survival.

## Discussion

First-generation, intratumourally-delivered CDN STING agonists, such as MK-1454 and ADU- S100, have consistently demonstrated immunostimulatory biological effects, but have largely failed to show sufficient clinical activity to warrant development beyond phase I/II trials (14). Recently, focus has switched to the development and clinical translation of second-generation STING agonists suitable for systemic delivery. diABZI, TAK-676, SR-717 and MSA-2 are non-CDN or CDN STING agonists that are active by systemic delivery through intravenous, subcutaneous and oral routes, respectively (15–18). Systemic administration of tumour cell- directed STING agonist antibody-drug conjugates has also been shown to exert anti-tumour activity through STING activation in tumour and myeloid cells in preclinical mouse models (19). These newly developed agents and administration modalities allow for thorough investigation of STING agonism as an approach of cancer immunotherapy, both preclinically and clinically.

Here, using preclinical models, we report immune- and anti-tumour effects of a novel, synthetic, systemically-delivered STING agonist, BI-1703880 (STINGa), an agent that is currently being tested in a phase I study (NCT05471856). Our data show that BI-1703880 is a potent and selective STING agonist *in vitro* and *ex vivo*, in both human and murine cells, and is systemically active *in vivo* in multiple mouse tumour models. BI-1703880 showed specific and dose-dependent activation of the STING signalling pathway and downstream immune activation. Enhanced immune effects and tumour growth control were seen when low-dose STINGa was combined with radiotherapy and this combination generated protective anti- tumour immune memory.

Our first step was to define the dose of STINGa that we should use in combination with RT. To study the effect of different doses of systemically administered BI-1703880, we used *in vivo* subcutaneous tumour models in immunocompetent mice and analysed plasma and intratumoural cytokine levels, tumour histology and dendritic cell activity in tumour-draining lymph nodes. The marked increase we observed in systemic interferon and cytokine levels following high-dose STINGa, combined with the knowledge that in a mouse model expressing an inducible constitutively active STING mutation (hSTING-N154S) exhibited rapid weight loss, hypercytokinemia and death (20), led us to assess multiple administrations of lower doses of STINGa. Given such concerns about high-dose STINGa and our desire to assess multiple administrations of the agent, we hypothesised that it would be safer to use repeated low-dose STINGa administration in combination with radiotherapy. Reassuringly, following low-dose intravenous STINGa delivery, the changes of cytokine levels in circulation were rapid and transient. Within 24 hours, all cytokine levels in blood, and most in tumours, had returned to basal levels. We elected to develop a twice-weekly low-dose STINGa schedule, positing that it would be beneficial for continued immune boosting. Transient induction of interferons in the TME supports T cell infiltration and dendritic cell activation, while prolonged or chronic exposure could suppress immunity and be detrimental (21). In our models, low-dose STINGa activated both dendritic cells and CD8+ T cells. Activation of both these cell types is essential for long-lasting antitumour activity (22, 23) and CD8 *in vivo* depletion experiments confirmed their importance in successful RT/STINGa combination therapy.

We hypothesised that delivering RT before STING agonist therapy would be optimal, facilitating an initial RT-induced increase in tumour antigenicity followed by STINGa-induced immune stimulatory adjuvant effects. In designing our studies, we were aware that such a schedule would deplete immune cells (especially highly radiosensitive T cells) from the tumour microenvironment, which might impair anti-tumour immunostimulation. Counterbalancing this effect, however, we speculated that those CD8+ T cells already residing in the tumour pre-treatment were either not antigen-reactive or exhausted and that depleting them with RT might promote repopulation of the tumour microenvironment with new, antigen-reactive T cells capable of initiating anti-tumour immune responses. To test this hypothesis, most of our analyses of the tumour microenvironment were performed ∼10-14 days after radiotherapy, to capture events relating to an ongoing active anti-tumour response. Our supposition was supported by previous studies that examined four different sequences of intratumoural DXMAA combined with radiotherapy. The combination that exerted the greatest tumour growth delay involved irradiation prior to administration of DMXAA (24).

The potential of radiotherapy to induce anti-tumour immune responses through cGAS/STING pathway activation has been extensively reviewed (7). Importantly, our data show that while both radiotherapy and low-dose STING agonist are suboptimal in controlling tumour growth as monotherapies, the combination yielded tumour growth control and cure. To study the interaction between radiotherapy and STINGa, we deliberately used a sub-therapeutic, short- course radiation dose (16 Gy in 2 fractions) which slows, but does not locally control, *in vivo* tumour growth in multiple mouse *in vivo* models. Treatment with a synthetic CDN derivative has previously improved the efficacy of radiotherapy in a murine pancreatic tumour model through initial TNFα-dependent haemorrhagic necrosis followed by CD8+ T cell-dependent tumour control (25). In contrast to our approach, in that study the STING ligand was injected intratumourally which probably contributed to the high degree of initial necrosis. Fractionated radiotherapy has also been shown to synergise with an inhaled nanoparticulate STING agonist in a model of lung metastasis (8). The authors reported activation of anti-tumour immunity in both irradiated and non-irradiated tumours, in line with the increase in CD8+ T cells with RT/STINGa combination therapy in our bi-flank model. Such results highlight the potential for generating systemic immune responses against multi-site disease following treatment with RT/STINGa.

Our selection of suboptimal doses of radiation and relatively low-dose STING agonist meant that there was room to further improve therapeutic efficacy in our tumour models. We hypothesised that this might be achieved through rational selection of immune checkpoint inhibitors (ICI) for combination therapy. To inform our choices, we performed bulk RNA sequencing to define the immune landscape of RT/STINGa-treated, as opposed to single agent-treated, tumours. These analyses revealed an increase in immune checkpoint transcripts (including Pdcd1 and Ctla4) with the combination of RT/STINGa. Subsequent flow cytometry confirmed that CD8+ T cells in tumours treated with combined RT/STINGa showed higher expression of exhaustion markers. PD-L1 expression is also known to be upregulated by several inflammatory cytokines and interferon signalling (26–28), and, in our experiments, STINGa induced expression of PD-L1 in tumours. Furthermore, our data generated from Nr4a3-tocky mice suggest that RT/STINGa treatment expands a subset of persistently TCR- signalling CD8+ T cells with an activated but potentially exhausted phenotype. Together, these findings indicated that additional immunotherapeutic intervention might enhance the efficacy of RT/STINGa. Targeting PD-1 and CTLA-4 offers the opportunity to exploit distinct mechanisms of action, potentially with different spatial and temporal effects. We targeted both PD-1 and CTLA-4 successfully to reverse RT/STINGa-induced T cell exhaustion and improve therapeutic efficacy. When comparing tumours treated with RT/STINGa to those treated with RT/STINGa/aPD-1/aCTLA-4 using bulk RNA sequencing, an increase in granzyme and perforin expression, most evidently in the TBPt-4C4 model, suggested an increase in cytotoxic cells in tumours receiving quadruple therapy. Taken together, our data suggests that RT/STINGa results in initial killing of tumour cells followed by recruitment of immune cells. Repeated STINGa administration enables priming and boosting DCs and the addition of aPD-1/aCTLA-4 expands the T cells, enhances their effector response and reinforces anti-tumour efficacy.

Interestingly, our experiments showed that the efficacy of RT/STINGa combination therapy depends on tumour size at the time of treatment initiation. While combined RT/STINGa therapy was sufficient to effect tumour clearance with smaller-sized tumours, the addition of aPD-1/aCTLA-4 ICI was needed to cure larger tumours. In addition to differences in tumour volume, other factors like vessel density, presence of hypoxia and immune and stromal cell composition might also impact the response to therapy and these factors are the subject of ongoing research in our laboratory. Both anti-CTLA-4 and anti-PD-1 checkpoint inhibitors, alone or in combination, have increased survival rates in patients with cancer in numerous studies (29). Clinical trials combining STING agonists and ICI are currently ongoing (14). The PD-1 pathway has a well-established role in regulating exhausted PD-1+ T cells. However, PD- 1 is also expressed on all T cells during activation and is known to be an important player in shaping both qualitative and quantitative aspects of CD8+ T cell effector and memory responses. Timing and duration of PD-1 blockade has shown to impact on CD8+ T cell memory generation after acute infection (30). In our studies, anti-PD-1 was given on the same days as STINGa, largely as an empirical and convenient approach. Further evaluation of the kinetics and amplitude of STINGa-induced upregulation of PD-L1 might allow us to refine scheduling of STINGa-ICI combinations. In addition, a deeper understanding of the spatial and temporal effects of RT and STINGa on T cell receptor signalling and T cell biology might enable further optimisation of the combination regiments. Such work is, at present, beyond the scope of our current studies.

STING agonists hold great but, not yet substantiated, promise for the treatment of solid cancers. Here, we show in preclinical mouse models that low doses of the novel BI-1703880 STINGa can function as a therapeutic combination partner to radiotherapy without signs of the toxicities associated with high dose STINGa dosing. Systemic delivery of STING agonists combined with locoregional radiotherapy opens a therapeutic window for enhancing cancer immunotherapy for either de novo immune responses in relatively non-inflamed tumours or for boosting pre-existing immune response in already inflamed tumours.

## Methods

### Compound

BI-1703880 STING agonist (STINGa) or vehicle were used for *in vivo* experiments. For *in vitro* and *ex vivo* assays, STINGa was dissolved in DMSO and added to the culture media. BI- 1703880 STINGa was provided by Boehringer-Ingelheim and is currently being evaluated in a clinical trial (clinicaltrials.gov NCT05471856).

### Human cells, cell lines and IRF reporter assay

The reporter cell line THP1-Blue^TM^ ISG (Invitrogen) were maintained according to the manufacturer’s instructions. Paired THP-1 cell lines were generated: parental (WT:wildtype STING; THP1-BlueISG-STINGwt) or STING-knockout (THP1-BlueISG-STINGKO). THP1-Blue ISG STING knockout (KO) cells were generated using the Crispr/CAS9 system with the ALL-IN construct U6gRNA-Cas9-2A-GFP-TMEM173 (Sigma #Target ID HS0000556293). To generate THP1 reporter cell lines for each of the five major STING isoforms expressed in humans, THP1- BlueISG-STING-KO cells were transduced with retrovirus containing each of the five different STING isoforms. Cell populations expressing graded expression of GFP were isolated by FACS and their STING protein expression level was evaluated by Western Blot analysis. Cell populations expressing STING to a similar extend as the endogenous STING level in the parental THP1-BlueISG cell line were selected for further experiments. To quantify STING activation the expression of the reporter gene SEAP (secreted embryonic alkaline phosphatase) was quantified by adding 90μL QUANTIBlue reagent, incubation of the samples at room temperature for 10-20 minutes and absorption was measured at reading the OD at 620nm. These cells allow the monitoring of STING-mediated IRF3 activation by determining the activity of SEAP.

Dendritic cells were derived from human peripheral blood mononuclear cells (PBMCs) from healthy donors obtained via the Austrian Red Cross using EasyStep Human Monocyte Isolate Kit (StemCell Technologies #19359) followed by cultivation of isolated cells in the presence of 1000 U/mL recombinant human GM-CSF (PeproTech #300-03) and 300 U/mL recombinant human IL-4 (PeproTech #200-04) for 5-7 days. Cells were treated STINGa (with 0-1-5-10μM) for 6 hours. Activation of the STING pathway was tested using Simple Western Immunoassays (ProteinSimple) according to the manufacturer’s protocol, using cell protein lysates from either THP-1 cells (wildtype or STING knockout) or from PBMCs from three different donors. The following antibodies were used: pSTING: CST #85735, STING: CST #13647, pTBK1: CST #5483, TBK1: CST #3504, pIRF3: Abcam #182859, IRF3: CST#11904 and GAPDH: Abcam 8245.

### Murine cell lines, tumour growth and *in vivo* treatments

Murine cell lines (mEER, TBPt-4C4, TBP-B79, TKP-4831, MC38, TC-1 and AT84) were cultured at 37°C in Dulbecco’s modified Eagle’s medium (DMEM) supplemented with 10% FBS (Gibco), 2.4mM L-glutamine, 60μg/mL penicillin, 100μg/mL streptomycin and 0.1mg/mL Primocin (Invivogen #ant-pm-1). DC2.4 cells were supplemented with β-Mercaptoethanol in culture. All cell lines were regularly tested for mycoplasma using MycoStrip^TM^ Mycoplasma Detection Kit (InvivoGen #rep-mysn). Cells were injected subcutaneously in the right flank of 7-10 week-old female mice. After cell inoculations, tumous were allowed to grow for 10-15 days depending on model and mice were randomised based on tumour volume before start of treatment. For orthotopic injections, 50μl of mEER cells were injected into the submucosal inner lip.

Wild-type C57Bl/6 or C3H females were purchased from Charles River Laboratories. NOD scid gamma (NSG) mice and C57Bl/6;Nr4a3 tocky mice were bred-in-house. All procedures involving animals were approved by the Animal Welfare and Ethical Review Board at The Institute of Cancer Research in accordance with National Home Office Regulations under the Animals (Scientific Procedures) Act 1986.

Mice in the STINGa group were given up to 6 systemically administered injections of STINGa, every 3-4 days, as 20 μmol/kg (high dose) or 7 μmol/kg (low dose). The first three injections were administered intravenously (i.v.) via the tail vein, and the subsequent two or three injections were administered intraperitoneally (i.p.). Systemic injections were switched to i.p. injections to refine compound administration to improve animal welfare due to technical limitations with multiple i.v. dosing to the tail vein. Bridging pharmacokinetics (PK) with i.p. dosing in mice were carried out to ensure comparability of the PK profiles with i.v. dosing, and resulted in i.v.-like PK profiles, with rapid absorption (Tmax at first sampling time point 15 minutes) and i.p. bioavailabilities of 37-99% in the dose range 2-20 μmol/kg. Vehicle solution was given systemically at the same days as STING agonist to all mice in vehicle or RT cohorts. Animals were irradiated under anaesthesia with Xylazine / Ketamine given by intraperitoneal injection. Local tumour radiotherapy was delivered in two fractions of 8Gy each on consecutive days using a AGO250 kV X-ray machine using lead shielding. Radiation dose was measured using a Farmer Chamber and Unidos-E Dosimeter (PTW). Mouse body-weights were measured twice weekly. Once palpable, tumours were measured using callipers twice-weekly and tumours volumes were calculated using the formula length x width x height (mm) x 0.5236, and data plotted in GraphPadPrism. Data from *in vivo* experiments are presented as mean +/- SEM. For survival experiments, tumours were allowed to reach 14 mm in any dimension and then sacrificed and censored. Kaplan-Meier survival curves were compared using the log-rank (Mantel-Cox) test to detect statistically significant differences between treatment groups.

For experiments involving MC38 tumours, the cells were sourced from Shanghai Ruilu Biotechnology co. LTD and wild-type C57Bl/6 females were purchased from Shanghai Lingchang Biotechnology Co., Ltd (Shanghai, China). Cells (1 million per mouse) were injected subcutaneously in the right flank of 6–10-week-old female mice. Mice were randomly allocated to 4 study groups (20 mice/group) when the mean tumour size reached 110 mm^3^. Local tumour radiotherapy was delivered in a single fraction of 8Gy using the X-RAD SmART device (Precision X-ray Inc) using lead shielding. STINGa was given intravenously via the tail vein at 7μmol/kg every 3 days at 4 occasions. Half of the mice per group (n=10 per time point) were euthanized 8 and 11 days post radiation-treatment, respectively, to collect tumours for further analysis.

*In vivo* antibodies were purchased from 2BScientific. *In vivo* depletion of CD8+ cells was performed using 200 μg of anti-CD8 (clone 2.43) or isotype control (LTF-2) injected via intraperitoneal injections every second day, starting at treatment day 0 (one day before RT). A total of 10 injections were given. Immune checkpoint inhibition was performed using 200 μg of anti-PD-1 (clone RMP1-14), 150 μg of anti-CTLA-4 (clone 9D9) or isotype controls (clones 2A3 and MPC-11) and were given using intraperitoneal injections two to three times per week for up to 8 injections.

### Cytokine analysis

For cytokine analysis from *in vitro*-treated THP-1 cells and human dendritic cells, they were seeded in 24-well plates, treated with STINGa for 24 hours at 37°C and cytokine levels were measured with 27- and 37-plex kits from Biorad. Data was measured with Bioplex3D and Bioplex200. For cytokine analysis of *in vivo* samples, mouse tumours or plasma were snap-frozen and analysed using Mouse ProcartaPlex Mix&Match 15-plex kit from Invitrogen (PPX- 15-MXFVNTG). Murine tumour cells were tested for STINGa induced IFNβ secretion using ELISA. Cells were seeded in 12-well plates at 250,000/well and treated the following day with DMSO, STINGa (5 mM and 10 mM) and reovirus (MOI10) for 48 hours. The media was collected following the previously described collection method and IFNβ was detected using the Mouse IFNbeta and Ancillary Reagent kits from R&D Systems (#DY8234-05 & #DY008B).

### Histology, immunohistochemistry and *in situ* hybridisation

Mouse tumours were formalin-fixed and paraffin-embedded, sectioned and stained with H&E or antibodies against CD8a (AbCam ab217344), CD4 (AbCam ab183685) or CD31 (Dianova DIA- 310). The staining was performed using the Agilent Autostainer Link48 instrument. CD8a and CD4 were detected using anti-rabbit EnVision polymer-HRP (Agilent K400311-2) and CD31 with anti-rat N-Histofine polymer-HRP (Nichirei 414311F) and demonstrated with DAB. The scanned images were analysed and scored using either QuPath software for bioimage analysis, the imaging analysis software Halo (Indica Labs) and/or by certified pathologists.

For Ifnβ *in situ* hybridisation (ISH), 4-µm-thick sections from FFPE tissue blocks of MC38 mouse tumours were stained with Leica RED-mIFNbeta RNAscope LS 2.5 Probe - Mm-Ifnb 1 (#406538). The ISH was carried out on the automated staining platform Leica Bond RX according to the manufacturer’s protocol. Evaluation was performed by a board-certified MD pathologist. A tissue classification algorithm was trained to divide histologic sections into: viable tumour, stroma, and necrosis. An imaging analysis algorithm was trained to recognise true positive IFNβ ISH signal. The amount of ISH signal was quantified as percent area of viable tumour (optical density).

### Western Blot analysis

Murine tumour cells were seeded in 6-well plates at 600,000/well and treated the following day with DMSO and STINGa (10 mM) for 1 hour. To harvest the cells were scraped into ice cold PBS and centrifuged at 2500xg for 2 mins at 4°C. The cells were resuspended and vortexed in radioimmunoprecipitation assay (RIPA) buffer (50 mM Tris (pH7.5), 150 mM, 1% NP40, 0.5% sodium deoxycholate and 0.1% SDS) supplemented with protease inhibitors (Roche #11836153001), 1 mM sodium orthovanadate and 1 mM sodium fluoride and lysed by snap freezing on dry ice then thawing on ice. Protein was quantified by using BCA protein assay reagent (Pierce #23225). 15mg of each sample was loaded and run on 10% NuPage Bis-Tris precast gel (Invitrogen #NP0302BOX) before transferring to polyvinylidene difluoride (PVDF) membrane (Thermo Scientific #88518). The membrane was blocked in 5% milk in TBS-Tween (0.1%) for 1 hour and incubated with primary antibodies overnight in at 4°C. Antibodies used were purchased from cell signalling: STING(D1V5L) (#50494), pSTING(Ser365) (#72971), IRF3(D83B9) (#4302), pIRF3(Ser396) (#29047) and Abcam: β-actin (#ab6276). Blots were developed with secondary anti-mouse or anti-rabbit antibodies conjugated to horseradish peroxidase (GE Healthcare #NA931 and GE Healthcare #NA934) and detected using the Super Signal chemiluminescent substrate (Pierce #34580) or Immobilon Western chemiluminescent HRP substrate (Millipore #WBKLS0500). Proteins were visualised using the Curex60 Image Processor (AGFA). MC38 western blot analysis was run using Simple Western Immunoassays (ProteinSimple) according to the manufacturer’s protocol and antibodies against pSTING: CST #72971, STING: CST #13647 and α-actinin: CST #3134. Untreated lysate from DC2.4 cells was used as positive control for STING protein expression.

### RNA extractions and bioinformatics analysis

Human THP-1 cells or human dendritic cells isolated from PBMCs were treated with STINGa (10 μM) for 6 hours *in vitro*. RNA was prepared using RNeasy Mini Kit (Qiagen) according to the manufacturer’s protocol and used for RNA sequencing and subsequent analysis. Mouse tumours were collected in RNAl*ater* RNA stabilisation reagent followed by homogenization in lysis buffer (Precellys 24 homogenizer, Bertin Technologies) and RNA isolated using RNeasy mini kit (Qiagen). RNA-sequencing libraries were prepared using the QuantSeq RNA Library Preparation Kit and subsequently sequenced on the NextSeq 500 using a single-end protocol with 75 cycles for experiments from TBPt-4C4, TBP-B79 and MC38 tumours, while a paired-end 100 cycles protocol was used for mEER tumours.

The STAR alignment software (v.2.7.6a) (31) was used to align reads to Ensembl Human genome version GRCh38.103 and Mouse reference genome version GRCm39.103. Aligned reads were further processed using HTSeq-count (HTSeq v0.12.4) (32) to quantify genomic features expression in each sample. Differential gene expression analysis was performed in R using the Bioconductor package DESeq2 (v1.42.0) (33). Lowly expressed genes as well as ribosomal genes and pseudogenes were excluded from the downstream analysis. Gene set variation analysis (GSVA) was performed using GSVA package (v1.50) (34) for hallmark gene sets (35) from the Molecular Signatures Database (MSigDB) accessed using msigdbr library (v7.5.1) (36). Heatmaps showing Z-scores of log2 transformed normalised counts and GSVA gene set scores were plotted using ComplexHeatmap library (v2.18.0) (37). Differential expression (DE) significance symbols (* for p-adj < 0.05, ** for p-adj < 0.01, *** for p-adj < 0.001 and **** for p-adj <0.0001) are marked in the heatmaps of respective experiment. Expression in the vehicle-treated group of each experiment was used as baseline for DE significance calculations (Wald test for gene expression and permutation test for GSEA; Benjamini and Hochberg method was used for multiple testing correction). Bulk RNA sequencing data and gene counts generated in this study have been deposited in the Gene Expression Omnibus (GEO) under accession number GSE295992.

### Flow cytometry analysis

Tumours and tumour-draining lymph nodes were dissected from experimental mice. Tumours were mechanically dissociated and enzymatically digested in PBS containing 0.5mg/mL collagenase VI, 0.2mg/mL DNase, 0.4mg/mL dispase and 4% trypsin for 30 min at 37°C on a shaker. Lymph nodes and digested tumours were smashed through 70 μm cell strainers and with FACS buffer (PBS containing 2% FBS and 1mM EDTA). Cells were centrifuged at 500xg for 5 mins at 4°C, pellets were resuspended in FACS buffer and plated in a U-bottom 96-well plate. The cells were centrifuged again at 800xg for 2 mins at 4°C and resuspended in Fixable Viability Dye eFluor 780 (ThermoFisher Scientific #65-0865-14) and Anti-Mouse CD16/CD32 (BD Biosciences #553142) for 15 mins at 4°C. Cells were then stained with extracellular stains for relevant antibodies, permeabilised using the FoxP3 Transcription Factor Staining Buffer Set (ThermoFisher Scientific #00-5523-00), intracellular stained and fixed with the IC Fixation Buffer (ThermoFisher Scientific #00-8222-49). Samples were analysed using the FACSymphony A5 (BD Bioscience). Acquired data was analysed using FlowJo and statistically significant differences between treatment groups were detected using one-way ANOVA or unpaired t tests, using GraphPadPrism.

Antibodies were obtained from BD Bioscience, eBioscience or Biolegend: aMHCI (clone: M1/42), aPD-L1 (clone: 10F.9G2), aCD45 (clone: 30-F11), aCD3 (clone: 17A2), aCD8 (clone: 53-6.6), aCD4 (clone: RM4-4), aCD25 (clone: PC61), aCD44 (clone: IM7), aOX-40 (clone: OX-86), aTIM-3 (clone: RMT3-23), a4-1BB (clone: RUO), aCD69 (clone: H1.2F3), aPD-1 (clone: 29F.A12), aTIGIT (clone: 1G9), aICOS (clone: C398.4A), aGITR (clone: DTA-1), aCD62L (clone: MEL-14), aFoxP3 (clone: FJK-16S), aKi67 (clone: 16A8), aCTLA-4 (clone: UC10-4B9), aGzmb (clone: GB11).

Tocky UMAP Analysis was performed largely as previously described (13). The threshold of Timer fluorescence was set using fully stained C57Bl/6 wildtype T cells from matched tissues. Timer data were then FSC-normalised and angle-transformed using algorithms which were previously reported (11). This allowed for the generation of a data frame that included both marker expression and Timer-related parameters (angle and intensity). The marker data used as input for UMAP were arcsinh-transformed to improve visualisation and extreme negative outliers were removed. UMAP was then applied to the dataset using the CRAN package *umap* (38) and the resulting two dimensional projection was visualised using heatmaps generated with the R package ggplot2 (39). After confirming the principal components of interest with PCA, k-means clustering was performed on those principal components. The optimal number of clusters was determined by Bayesian Information Criterion (BIC) using the R *mclust* package (40). Each cluster was then projected back onto the UMAP space, enabling visualisation of both cluster boundaries and their frequency distribution across samples. Finally, maker and Timer fluorescence patterns within individual clusters were evaluated in R and FlowJo (v.10).

## Acknowledgements

The work presented in this article has been generated within a research collaboration with Boehringer Ingelheim. We would like to acknowledge the following Boehringer Ingelheim team members: Susy Straubinger (technical assistance), Otmar Schaaf and Ida Dinold (formulation of STINGa compound), Luciene Borges (MC38 flow myeloid panel design) and Francesca Trapani (histology discussions). We also thank Eva Crespo-Rodriguez, Bahire Kalfaoglu, Martin McLaughlin and Ioannis Roxanis, from the Institute of Cancer Research, for technical support, support in data analyses and histology discussions. We thank Harriet Whittock, Peter John-Baptiste and team members of the Biological Service Unit, the Flow Cytometry facility and Breast Cancer Now histopathology core facility at the Institute of Cancer Research for technical assistance.

This work was funded by Boehringer Ingelheim and further supported by research grants from Anthony Long Charitable Trust, Archobaleno Cancer Trust, The International Centre for Recurrent Head and Neck Cancer, RadNet 2 (RRCOER-Jun24/100006) and Cancer Research UK (DRCRPG-Nov22/100008).

## Abbreviations

APC: antigen-presenting cell
cDC: conventional dendritic cell
CDN: cyclic dinucleotide
cGAS: cyclic GMP-AMP synthase
CRS: cytokine release syndrome
CTLA-4: cytotoxic T-lymphocyte-associated protein 4
DC: dendritic cell
ICI: immune checkpoint inhibitor
IFN: interferon
Inj.: injection
i.p.: intraperitoneal
i.v.: intravenous
ISH: *in situ* hybridisation
IRF3: interferon-regulatory factor 3
KO: knockout
NFκB: nuclear factor kappa B
NSG: NOD scid gamma
PK: pharmacokinetics
PD-1: programmed death-1
PD-L1: programmed death-1 ligand
RT: radiotherapy
STING: stimulator of interferon genes
STINGa: STING agonist
TBK1: TANK-binding kinase 1
TCR: T cell receptor
TME: tumour microenvironment
WT: wildtype

## Supplementary Figure Legends

**Supplementary figure 1.**
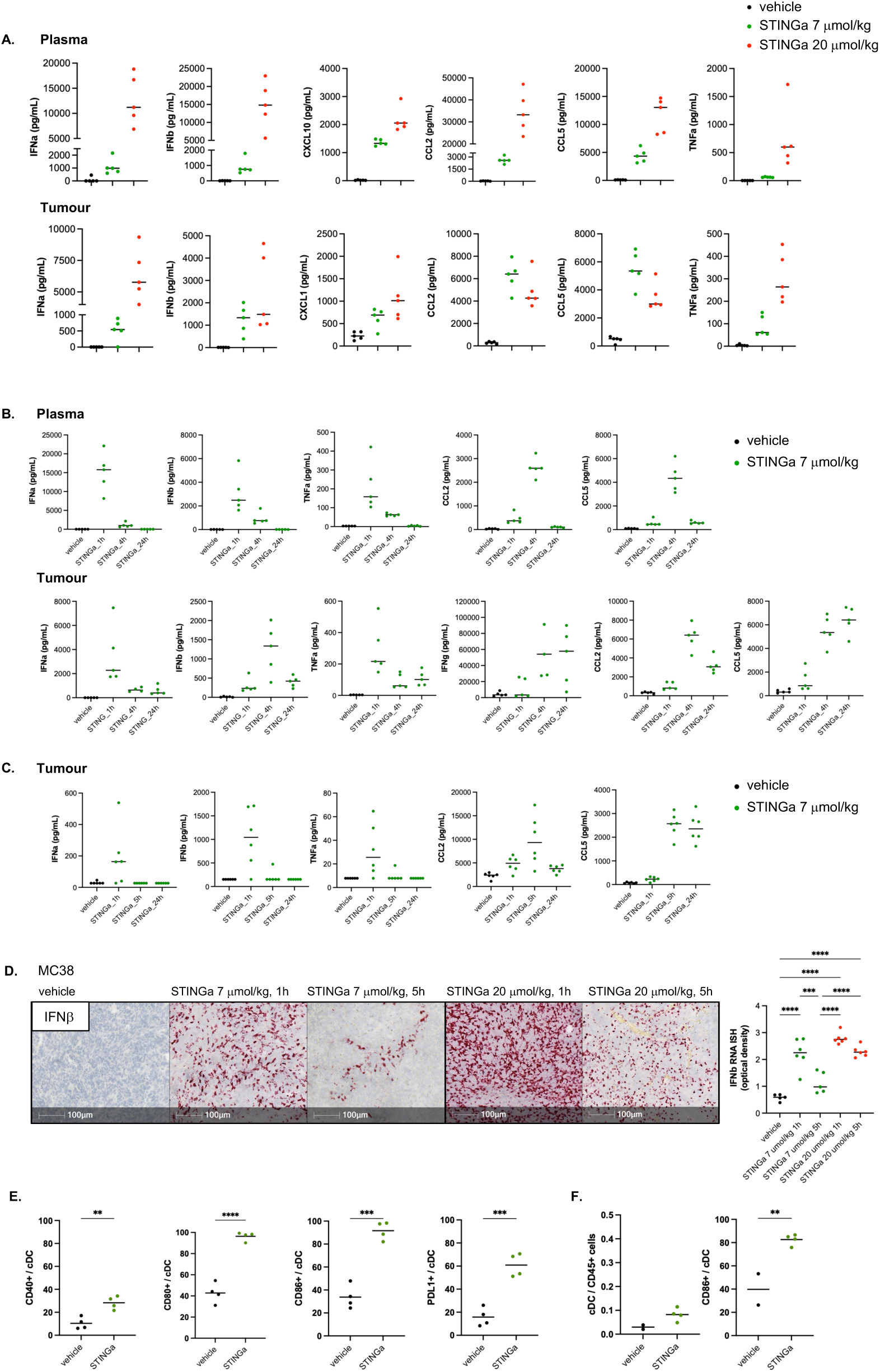
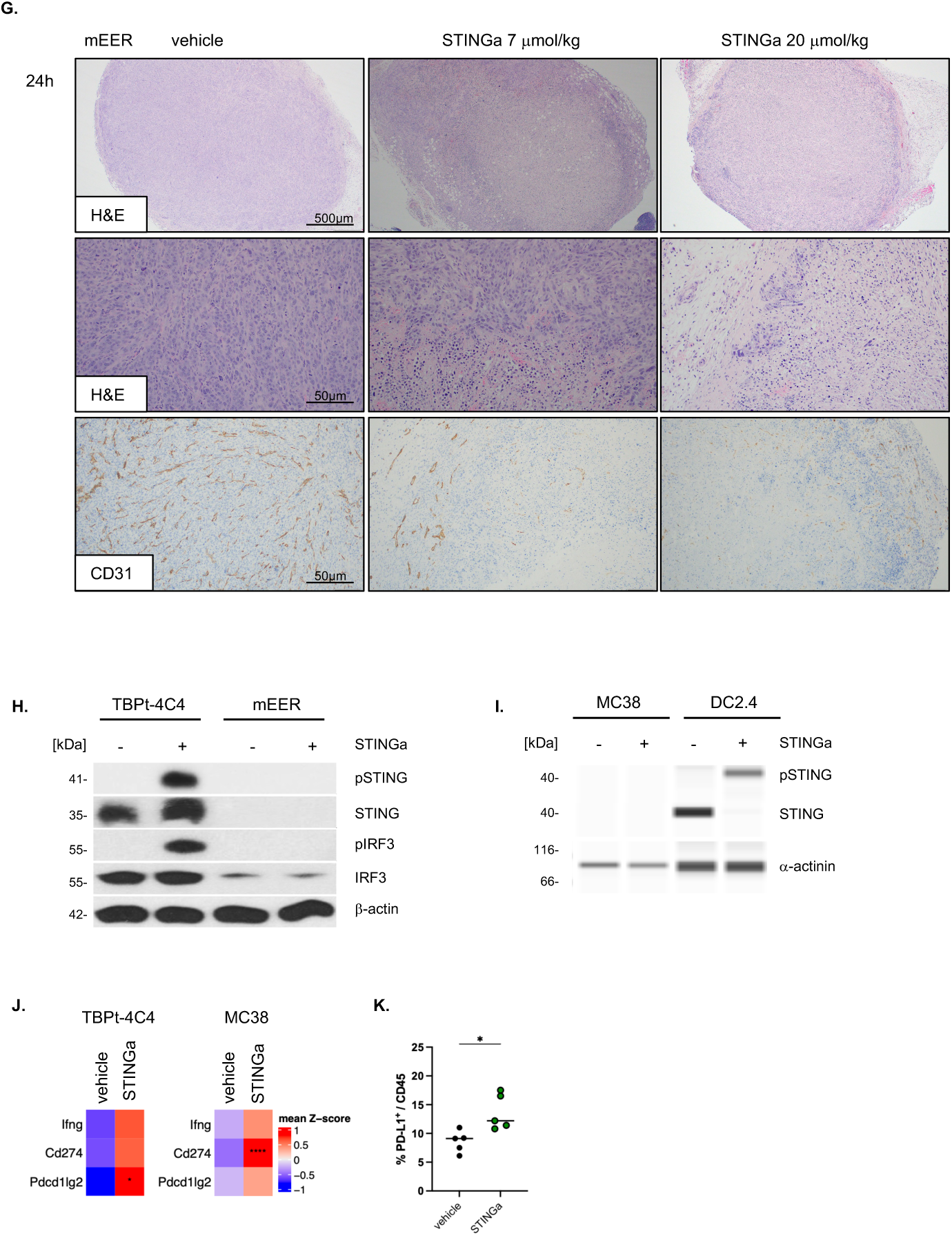
The effect of STING agonist on plasma and tumour cytokine levels, and tumour histology is dose- and time dependent. C57Bl/6 mice were subcutaneously injected with either mEER or MC38 tumour cells and treated systemically by intravenous injections with 7 mmol/kg or 20 mmol/kg of STING agonist. **(A)** Cytokine levels in blood and mEER tumours, 4 hours after a single dose of intravenous STING agonist were analysed. **(B)** Cytokine levels in plasma and mEER tumours, 1, 4 and 24 hours after a single dose of intravenous STING agonist (7 mmol/kg). **(C)** Cytokine levels in MC38 tumours, 1, 5 and 24 hours after a single dose of intravenous STING agonist (7 mmol/kg). **(D)** IFNb mRNA was analysed in MC38 tumours using *in situ* hybridisation (ISH), 1 or 5 hours after a single dose of STINGa. Optical density values were analysed and plotted to the right. One way ANOVA was performed to test for statistically significant differences between groups. **(E)** Tumour-draining lymph nodes (tdLN) from mEER tumour-bearing mice were analysed using flow cytometry 24 hours following one injection of STING agonist (7 mmol/kg). Percentage of CD40+ / CD80+ / CD86+ or PD-L1+ cDCs were plotted and unpaired t-tests performed. **(F)** Tumour-draining lymph nodes (tdLN) from MC38 tumour-bearing mice were analysed following injection of STING agonist (7 mmol/kg) for percentage of cDCs / CD45 cells and percentage of CD86+ cDCs using flow cytometry. Unpaired t-test was performed to test for statistically significant differences between treatment groups. **(G)** Representative images of H&E and CD31- stained photomicrographs of mEER tumours treated with STING agonist 7 mmol/kg or 20 mmol/kg *in vivo* for 24 hours. FFPE tumour sections were analysed. Scale bars: 500mm and 50mm. **(H)** TBPt-4C4 and mEER tumour cell lines were treated with STINGa (10mM, 1 hour) and subsequently lysed and analysed for STING pathway activation using Western Blotting. Antibodies against phospho-STING, STING, phospho-IRF3 and IRF3 were used. b-actin served as loading control. **(I)** MC38 cells were treated with STINGa (10mM, 6 hours) and subsequently lysed and analysed for STING pathway activation using Western Blotting and antibodies against pSTING and STING. a-actinin was used as loading control and lysates from STINGa treated DC2.4 cells served as positive control. **(J)** Ifng, PD-L1 and PD-L2 transcripts were analysed from RNAseq analysis of bulk TBPt-4C4 (vehicle: n=4 and STINGa: n=5) and MC38 (vehicle: n=10 and STINGa: n=6) tumours. Tumour-bearing mice received 4 injections of 7 mmol/kg STINGa or vehicle and tumours were collected 24 hours post the last STINGa injection. **(K)** TBPt-4C4 tumours were analysed for PD-L1+ - CD45+ cells using flow cytometry 24 hours after a single injection of 7 mmol/kg STINGa.

**Supplementary figure 2.**
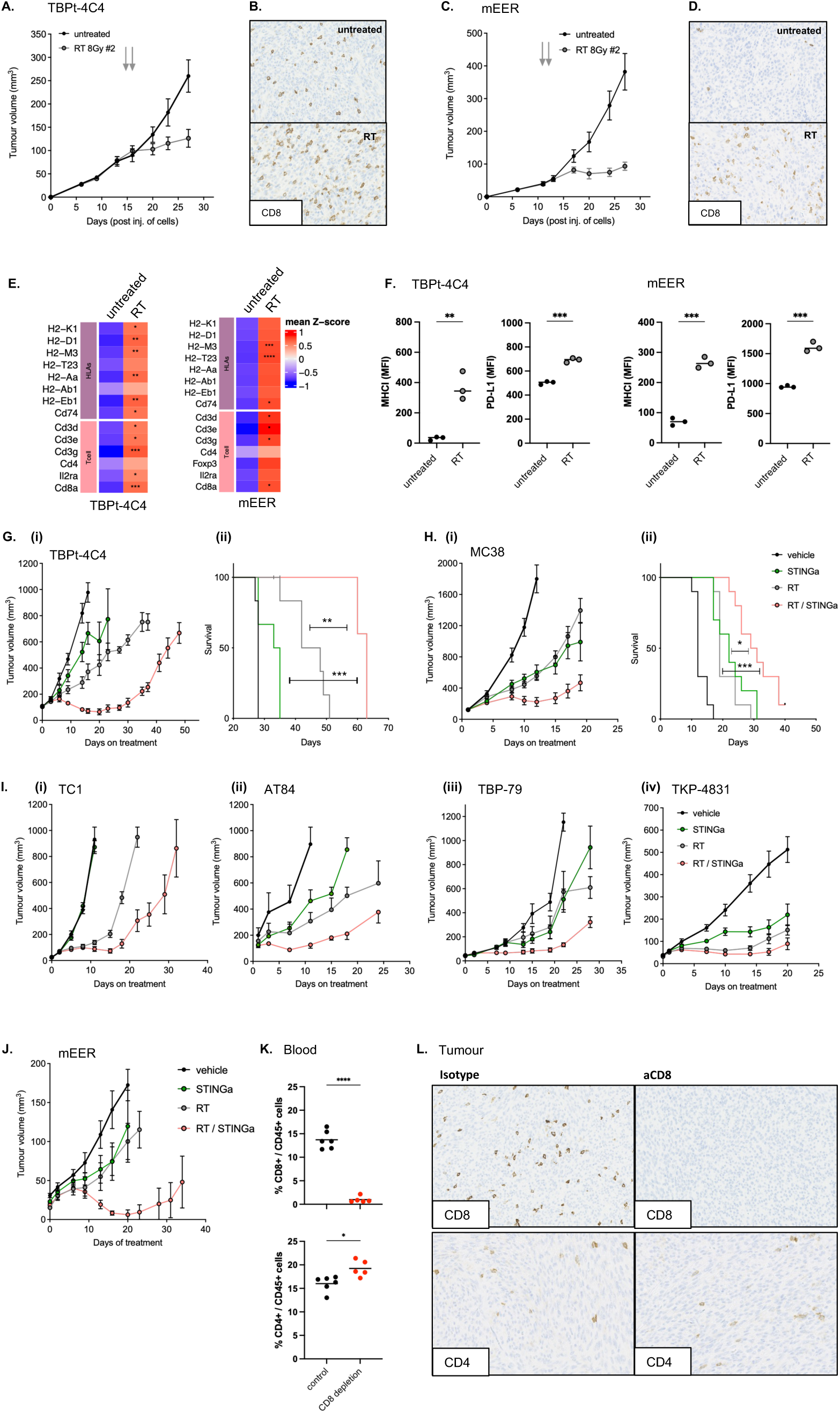
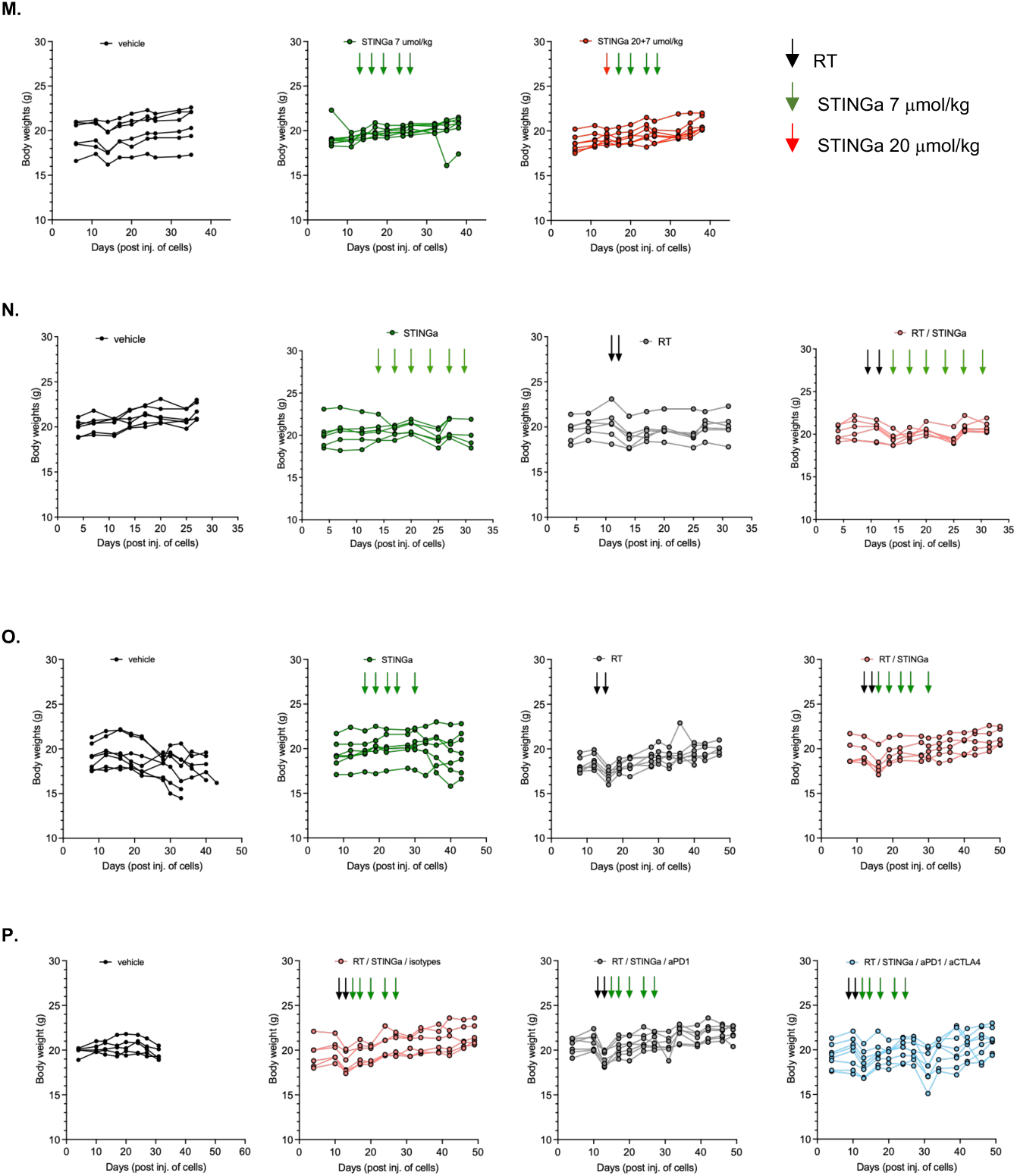
The combination of radiotherapy and STING agonist improves therapeutic outcome and is dependent on a functional immune system. C57Bl/6 or C3H mice were subcutaneously or orthotopically (lip) injected with tumour cells as indicated and treated with 2 fractions of 8Gy radiotherapy (RT) on consecutive days followed by 4-6 systemic injections (inj.) of STING agonist at 7 mmol/kg, every 3-4 days. **(A)** TBPt-4C4 tumour growth graph following 2 fractions of 8Gy RT. **(B)** Representative CD8+ staining of TBPt-4C4 tumours 14 days after RT. **(C)** mEER tumour growth graph following 2 fractions of 8Gy RT. **(D)** Representative CD8+ staining of mEER tumours 10 days after RT. **(E)** Bulk RNAseq was performed on tumours 14 days after start of radiotherapy (n=3-5 tumours / group) and analysed for HLA and T cell transcripts. (F) MHCI and PD-L1 staining (MFI) on live tumours cells (TBPt-4C4 and mEER) treated *in vitro* with 2 fractions of 8Gy RT and analysed 24 hours later using flow cytometry. **(G)** Tumour growth **(i)** and survival **(ii)** graphs of mice treated with radiotherapy and STING agonist (6 inj.) in the TBPt-4C4 subcutaneous tumour model, n=6-7 mice / group. Log-rank (Mantel-Cox) test was performed on Kaplan-Meier survival graphs: vehicle vs RT/STINGa: P value 0.0006 (***), RT vs RT/STINGa: P value 0.0011 (**), STINGa vs RT/STINGa: P value 0.0007 (***). **(H)** Tumour growth **(i)** and survival **(ii)** graphs of mice treated with radiotherapy (1 fraction of 8Gy) and STING agonist (4 inj.) in the MC38 subcutaneous tumour model, n=10 mice / group. Log-rank (Mantel-Cox) test was performed on Kaplan-Meier survival graphs: vehicle vs RT/STINGa: P value <0.0001 (****), RT vs RT/STINGa: P value 0.0007 (***), STINGa vs RT/STINGa: P value 0.0105 (*). **(I)** Tumour growth graphs of mice treated with 2 fractions of 8Gy RT and STING agonist (6 inj.) in the TC1 **(i)**, or (5 inj.) in the AT84 **(ii)**, TBP-B79 **(iii)** and TKP-4831 **(iv)** subcutaneous tumour models, n=6 mice / group. **(J)** Tumour growth graph of mice treated with 2 fractions of 6Gy RT and STING agonist (4 inj.) in the mEER orthotopic lip model, n=4-6 mice / group. **(K)** Flow cytometry analysis of CD4+ and CD8+ T cells in blood from mice treated with isotype control or aCD8 depleting antibody. **(L)** IHC analysis of CD8 and CD4 stained TBPt-4C4 tumours treated with isotype control or a CD8 depleting antibody. **(M)** C57Bl/6 mice were subcutaneously injected with TBPt-4C4 tumour cells and treated with STING agonist at 7 mmol/kg in 5 injections or one dose of 20 mmol/kg followed by 4 doses of 7 mmol/kg. Individual body weights over time from mice on treatment, corresponding to the experiment presented in Figure 2A. **(N)** C57Bl/6 mice were subcutaneously injected with TBPt-4C4 tumour cells and treated with 2 fractions of 8Gy RT on consecutive days followed by 5 systemic injections of STING agonist at 7 mmol/kg. Individual body weights over time from mice on treatment, corresponding to the experiment presented in Supplementary figure 3E. **(O)** C57Bl/6 mice were subcutaneously injected with mEER tumour cells and treated with 2 fractions of 8Gy radiotherapy on consecutive days followed by 5 systemic injections of STING agonist at 7 mmol/kg. Individual body weights over time from mice on treatment, corresponding to the experiment presented in Figure 3C. **(P)** C57Bl/6 mice were subcutaneously injected with TBPt-4C4 tumour cells and treated with 2 fractions of 8Gy RT on consecutive days followed by 5 systemic injections of STING agonist at 7 mmol/kg and immune checkpoint inhibitors. Individual body weights over time from mice on treatment, corresponding to the experiment presented in Supplementary figure 5B. Vehicle solution was given at the same days as STING agonist to all mice in vehicle or RT cohorts.

**Supplementary figure 3.**
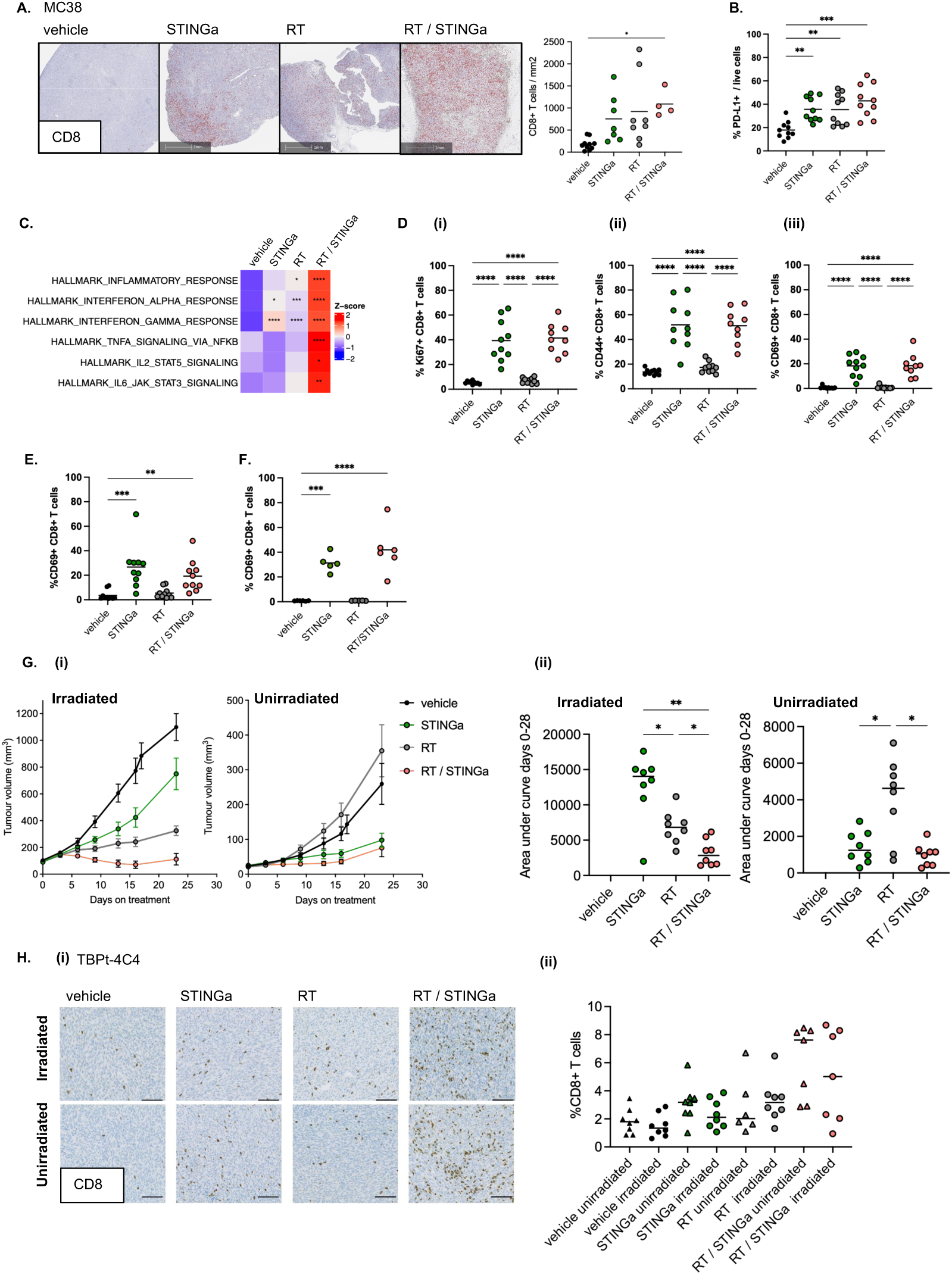
The combination of radiotherapy and STING agonist increases activated CD8+ T cells in tumours and in tumour-draining lymph nodes. **(A)** Immunohistochemistry images of representative MC38 tumours stained for CD8 following treatment with radiotherapy (1 fraction of 8Gy) and STING agonist. Quantification of CD8+ T cells in the tumours were analysed using Halo and plotted to the right. **(B)** TBPt-4C4 tumour- bearing mice were treated as in Figure 1A and all live cells in tumours were analysed for PD- L1 expression using flow cytometry 7 days after start of radiotherapy. **(C)** MC38 tumour- bearing mice were treated with RT and STING agonist and bulk RNAseq was performed on tumours 9 days after start of radiotherapy treatment and hallmark pathway analysis performed (n= 10 (vehicle), 9 (STINGa), 10 (RT) and 3 (RT/STINGa) tumours / group). Comparison of all groups is versus vehicle. **(D)** Mice bearing MC38 tumours were treated with RT and STING agonist and Ki67 **(i)**, CD44 **(ii)** and CD69 **(iii)** expressing CD8+ T cells in tumours were analysed using flow cytometry 11 days after radiotherapy. **(E)** TBPt-4C4 tumour-bearing mice were treated as in Figure 1A and CD8+ T cells in tumour-draining lymph nodes were analysed for CD69 expression using flow cytometry 7 days after start of radiotherapy. **(F)** mEER tumour-bearing mice were treated as in Figure 1A and CD8+ T cells in tumour-draining lymph nodes were analysed for CD69 expression using flow cytometry 10 days after start of radiotherapy. One way ANOVA was performed to test for statistically significant between treatment groups. **(G)** Mice bearing TBPt-4C4 tumours on both flanks were treated with radiotherapy on one flank and STING agonist systemically (5 inj. of 7 mmol/kg), n=8 mice / group. Tumour growth graphs **(i)** and area under curve analysis for day 0-28 **(ii)**. **(H)** CD8 stained immunohistochemistry images of representative TBPt-4C4 tumours at the end of therapy experiment (irradiated and unirradiated tumour of the same mouse), following treatment with radiotherapy and STING agonist **(i)**. Scale bar 100mm. Quantification of CD8+ T cells in the tumours were analysed using QuPath **(ii)**.

**Supplementary figure 4.**
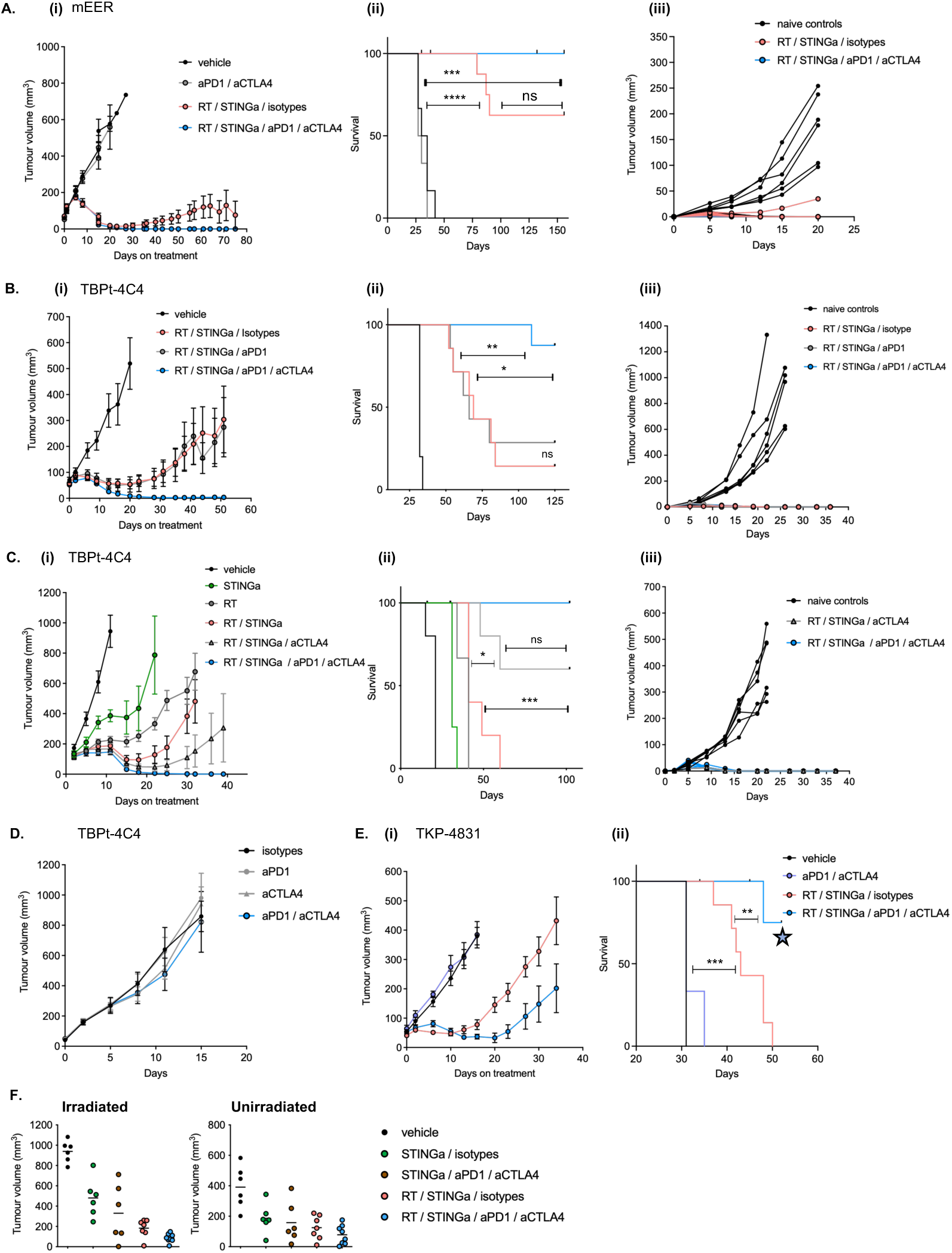
The addition of immune checkpoint inhibitors aPD-1 / aCTLA-4 improves therapeutic outcome. Tumour-bearing mice were treated with 2 fractions of 8Gy radiotherapy (RT) on consecutive days followed by 5 systemic injections (inj.) of STING agonist, every 3-4 days, and the combination immune checkpoint inhibitors aPD-1 / aCTLA-4. **(A)** Tumour growth **(i)**, survival **(ii)** and rechallenge **(iii**) graphs of the mEER subcutaneous model, n=6-8 mice / group. 5 out of 8 mice were cured of tumour in the RT / STINGa / isotype group and 6 out of 6 mice were cured of tumour in the RT / STINGa / aPD-1 / aCTLA-4 group. Log-rank (Mantel-Cox) test was performed on Kaplan-Meier survival graphs: vehicle vs RT/STINGa/isotypes: P value <0.0001 (****), vehicle vs RT/STINGa/aPD-1/aCTLA-4: P value 0.0002 (***), RT/STINGa/isotypes vs RT/STINGa/aPD-1/aCTLA-4: P value 0.1069 (ns). **(B)** Tumour growth **(i)**, survival **(ii)** and rechallenge **(iii)** graphs of the TBPt-4C4 subcutaneous model, n=6-7 mice / group. 1 out of 7 mice were cured of tumour in the RT / STINGa / isotype group, 2 out of 7 were cured in the RT / STINGa / aPD1 group and 7 out of 8 in the RT / STINGa / aPD-1 / aCTLA-4 group. Log-rank (Mantel-Cox) test was performed on Kaplan-Meier survival graphs: RT/STINGa/isotypes vs RT/STINGa/aPD-1: P value 0.8744 (ns). RT/STINGa/isotypes vs RT/STINGa/aPD-1-aCTLA-4: P value 0.0014 (**). RT/STINGa/aPD-1 vs RT/STINGa/aPD-1-aCTLA-4: P value 0.0109 (*). **(C)** Tumour growth **(i)**, survival **(ii)** and rechallenge **(iii)** graphs of the TBPt-4C4 subcutaneous model, n=6-7 mice / group. In this experiment STINGa (6 inj.) was administered at a dose of 7-13umol/kg. 0 out of 6 mice were cured of tumour in the RT / STINGa / isotype group, 2 out of 7 were cured in the RT / STINGa / aCTLA-4 group and 6 out of 7 in the RT / STINGa / aPD-1 / aCTLA-4 group. Log-rank (Mantel-Cox) test was performed on Kaplan-Meier survival graphs: RT/STINGa/isotypes vs RT/STINGa/aCTLA-4: P value 0.0265 (*). RT/STINGa/isotypes vs RT/STINGa/aPD-1-aCTLA-4: P value 0.0003 (***). RT/STINGa/aCTLA-4 vs RT/STINGa/aPD-1-aCTLA-4: P value 0.0766 (ns). **(D)** Tumour growth graph of mice treated with control single- agent immune checkpoint inhibitors aPD-1, aCTLA- 4 or the combination of aPD-1 / aCTLA-4 in the TBPt-4C4 subcutaneous model, n=5 mice / group. **(E)** Tumour growth **(i)** and survival **(ii)** graphs of the TKP-4831 subcutaneous model, n=7-8 mice / group. Log-rank (Mantel-Cox) test was performed on Kaplan-Meier survival graphs: vehicle vs RT/STINGa/isotypes: P value 0.0009 (***), RT/STINGa/isotypes vs RT/STINGa/aPD-1/aCTLA-4: P value 0.0086 (**). Blue star: the experiment was ended at day 52. **(F)** Mice bearing bi-flank TBPt-4C4 tumours were treated as in Figure 1A with the addition of aPD-1 / aCTLA-4 on the same days as STING agonist, tumour volumes of right flank (irradiated) and left flank (unirradiated) tumours were plotted on day 14 after start of radiotherapy.

